# Precise cell recovery by cell nucleus united transcript (CellCUT) for enhanced spatial transcriptomics

**DOI:** 10.1101/2024.05.28.596350

**Authors:** Bei Hong, Bo Zeng, Huimin Feng, Zeyuan Liu, Qi Ni, Wei Wang, Mayuqing Li, Meng Yang, Mengdi Wang, Le Sun, Suijuan Zhong, Qian Wu, Xiaoqun Wang

## Abstract

Cell segmentation is the first step in parsing spatial transcriptomic data, often a challenging task. Existing cell segmentation methods do not fully leverage spatial cues between nuclear images and transcripts, tending to produce undesirable cell profiles for densely packed cells. Here, we propose CellCUT to perform cell segmentation and transcript assignment without additional manual annotations. CellCUT provides a flexible computational framework that maintains high segmentation accuracy across diverse tissues and spatial transcriptomics protocols, showing superior capabilities compared to state-of-the-art methods. CellCUT is a robust model to deal with undesirable data such as low contrast intensity, localized absence of transcripts, and blurred images. CellCUT supports a human-in-the-loop workflow to enhance its generalizability to customized datasets. CellCUT identifies subcellular structures, enabling insights at both the single-cell and subcellular levels.

## Introduction

The emergence of spatial transcriptomic (ST) technologies, with its remarkable ability to reveal the spatial location information of individual cells, has ushered in a new era of understanding tissue structures and functions^1, 2^. Various ST technologies that achieve single-cell resolution for different genes have emerged, and they can be divided into two major categories^3^: imaging-based ST technology and sequencing-based ST technology. Typically, imaging-based ST technology provides high-resolution single-cell spatial data, has become one of the popularly used protocol. It can be further subdivided into in situ sequencing (ISS)-based methods (e.g., STARmap^4^, NanoString CosMx SMI^5^, 10x Genomics Xenium^6^, and etc.) and in situ hybridization (ISH)-based methods (e.g., MERFISH^7^, seqFISH^8^, seqFISH+^9^, and etc.). The rapid development trends of ST technologies vary in terms of their technical parameters and typically involve tradeoffs among the spatial resolution, the number of genes profiled and accuracy, posing new challenges for the subsequent data analysis^10^.

Accurately extracting single-cell gene expression profiles from spatially resolved data underpins downstream analyses and biological interpretations. This process typically involves two critical steps^11^: cell segmentation to delineate individual cell boundaries and RNA molecules assignment to each identified cell. Delineating cell boundaries within densely packed, overlapping structures with uneven illumination and low signal-to-noise ratios presents a challenging task due to similar textures and indistinct borders. Second, even with the excellent performance demonstrated by deep learning models, collecting manually annotated data for model training is very time-consuming and tedious. Furthermore, erroneous or absence information of local clues may be introduced during sample preparation, affecting the segmentation accuracy. Therefore, providing an efficient and accurate computational framework for upstream tasks has emerged as a key technology hotspot in field.

Efforts have been made toward the development of cell segmentation algorithms in ST, which can be broadly categorized into two groups: (1) image-based methods, where cells are recovered from staining images by image processing, and (2) molecule-based methods, which attempt to identify individual cells by clustering RNA molecules. The former methods include traditional image processing, e.g., Watershed^12^, or deep learning, e.g., Cellpose^13^, Stardist^14^, DeepCell^15^, GeneSegNet^16^, JSTA^17^, and etc. However, these methods either do not fully consider molecular spatial information or struggle with image quality, such as weak staining, overlapping cells, and blurred images. The later approaches, including pciSeq^18^, ClusterMap^19^, Baysor^20^, and SCS^21^, attempt to determine individual cells by clustering RNA molecules, and all transcripts in each cluster are assigned to a single cell. However, such methods often struggle when the given gene distribution is homogeneous, and they are sensitive to the number of gene throughputs or initial cell segmentation priors. In addition, several methods attempt to optimize their results with additional cell type priors^17, 22, 23^. Both of these approaches suffer from potential limitations in regions with low imaging resolution or densely packed cells, as relying solely on images or transcripts may not provide sufficient local clues to achieve satisfactory cell segmentation results.

Here, we propose CellCUT (**Cell** recovery by **C**ell nucleus **U**nited **T**ranscript), an innovative and flexible computational framework to jointly optimize cell segmentation and transcript assignment for ST data, to fill in the aforementioned gaps. There are three key properties of CellCUT. First, it performs cell segmentation robustly and precisely performance without requiring additional manual annotations for model training. Second, it applies a graph neural network (GNN) to estimate the inherent affinity between gene molecules, which enables the capture of the cytoplasmic region. Third, it builds a constrained graph partitioning model for nuclear and transcript information to solve the two associated complementary optimization tasks, which does not need to specify the clusters number, and allows to identify the subcellular components. With the above properties, CellCUT not only compensates for the lack of information contained in individual data sources, but also greatly improves the cell segmentation accuracy for ST data. We apply CellCUT to nine ST datasets derived from a range of tissues and various platforms, and demonstrate that CellCUT is superior to eight competitive algorithms including Watershed, Cellpose, Stardist, Baysor, ClusterMap, and SCS, in terms of cell counts, the number of transcripts per cell, and cell border estimation. We use CellCUT to analyze subcellular RNA localization, and the results agree with previous knowledge, further validating the accuracy of our segmentation approach. We show that the accurate cell segmentation of CellCUT can reveal the finer spatial structures compared to competitive algorithms.

## Results

### Overview of CellCUT

In this section, we present a concise overview of CellCUT. A detailed description of the algorithm can be found in the “Methods” section. Briefly, CellCUT takes staining images and decoded transcripts as its inputs, and simultaneously outputs single-cell instance segmentation results as well as transcript assignments to each cell (Fig. 1a). The schematic diagram of CellCUT is shown in Fig. 1b. Specifically, to fully leverage spatial consensus cues, CellCUT consists of three main parts: a robust pretrained pixel embedding network for images (Fig. 1b-Ⅰ), a generalizable graph embedding network for transcripts (Fig. 1b-Ⅱ&Ⅲ), and a flexible graph partitioning model for information integration (Fig. 1b-Ⅳ).

**Fig. 1.**
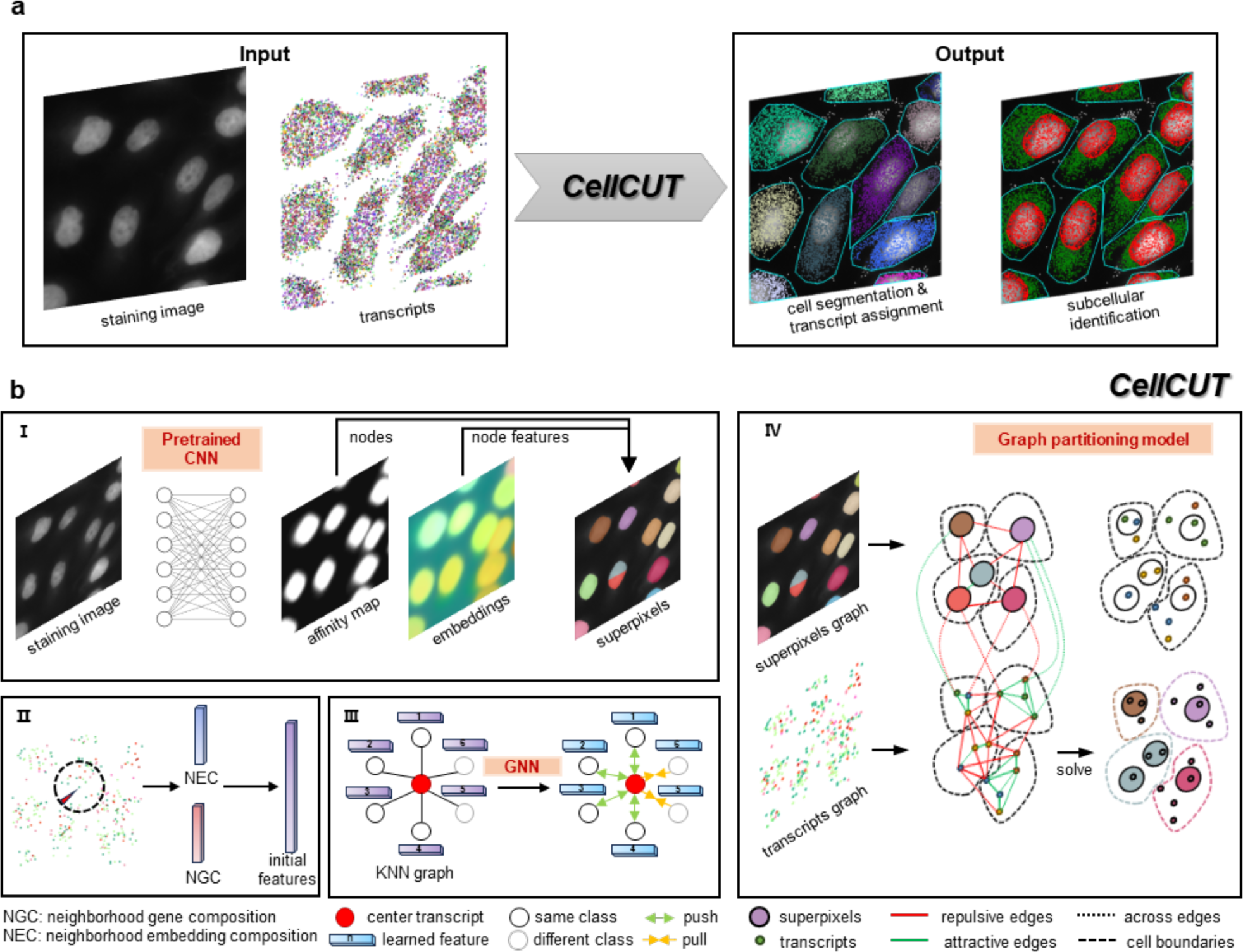
Schematic diagram and cell segmentation performances of CellCUT. **a**, Taking the nucleus staining image and molecule spatial information as input, CellCUT outputs the cell segmentation and transcript assignment results simultaneously. **b**, CellCUT includes four steps. Ⅰ, a neural network is trained on nucleus staining images to predict an affinity map as well as the pixel embeddings, and these two outputs are combined into a graph, where the nodes are superpixels of the nucleus and the edges denote similarities. Ⅱ, the neighborhood gene composition (NGC) and neighborhood embedding composition (NEC) are extracted for each transcript. Ⅲ, a GAT model is trained on the extracted feature vectors to learn the embeddings of adjacent transcripts. Ⅳ, combining superpixels graph and transcripts graph to obtain cell segmentation and transcript assignment results via a graph partitioning model.

First, for staining images, CellCUT learns two outputs via a convolutional neural network (CNN), one is the probability that each pixel belongs to a cell, and the other is the high-dimensional embeddings for each pixel (Fig. 1b-Ⅰ). The goal of embeddings is for pixels belonging to the same cell to be close to each other, and for pixels belonging to the different cell to be far away from each other. To extend the learning capability of CellCUT, a pretrained network model based on extensive publicly available nucleus datasets (Method) is trained to obtain robust and generalized nucleus features.

Second, for each transcript, the neighborhood gene composition (NGC) and neighborhood embedding composition (NEC) are first extracted as the initial feature vectors (Fig. 1b-Ⅱ), where NGC denotes the neighboring gene composition and NEC denotes the neighboring pixel embedding composition (Methods). A k-nearest neighbor (KNN) graph among the transcripts is constructed based on Euclidean distances, and the feature vectors are updated by using a graph attention network (GAT) to encode both gene expression profiles and embedded spatial information (Fig. 1b-Ⅲ). The goal of the GAT is for the learned feature vectors of transcripts belonging to the same cell to be close to each other, while the learned feature vectors of transcripts belonging to different cells should be far from each other (Fig. 1b-Ⅲ).

Third, a joint optimization graph partitioning model for nucleus superpixels and transcripts is constructed based on image pixel embeddings and the learned transcript features (Fig. 1b-Ⅳ). This model models the similarity of different nodes by edge weights derived from embeddings (Methods), and introduces the across edges to combine two kinds of information. The final cell segmentation and transcript assignment results are obtained by solving the graph model. For visualization, cell boundaries are obtained by convex hulls belonging to the same category of nodes. By integrating these two sources of information, CellCUT achieves precise transcript assignment and boundary recovery at the single-cell and subcellular level.

### CellCUT provides accurate cell segmentation and transcript assignment performance

Cellular morphology exhibits substantial variability across different tissues, falling into two main categories: cells with densely packed nuclei and small cytoplasmic regions, or sparsely distributed cells with large cytoplasmic regions. To comprehensively evaluate different cell segmentation methods, we evaluated their performance on five high-resolution datasets from the above two perspectives cover a wide range of platforms, imaging resolutions, and gene throughput (Fig. 2, Supplementary Fig. S1-S4), including seqFISH+ NIH/3T3 dataset^9^, MERFISH u2os dataset^7^, NanoString non-small-cell lung cancer (NSCLC) dataset^5^, Xenium human breast cancer dataset^6^, ISS CA1 dataset^18^ (Fig. 2a). To benchmark the study, we compared CellCUT with Watershed and two popular segmentation methods, named Cellpose and Stardist, which were based on deep learning. Appropriate pretrained models of the deep learning methods were used for evaluation purposes (Methods). Additionally, we compared CellCUT with two representative molecule-based methods, named ClusterMap and Baysor, both with and without using nucleus priors. We performed a comprehensive analysis using multiple metrics, including cell counts (Fig. 2b), average number of transcripts per cell (Fig. 2c), and adjusted mutual information (AMI) compared to the manual annotations (Fig. 2e).

**Fig. 2.**
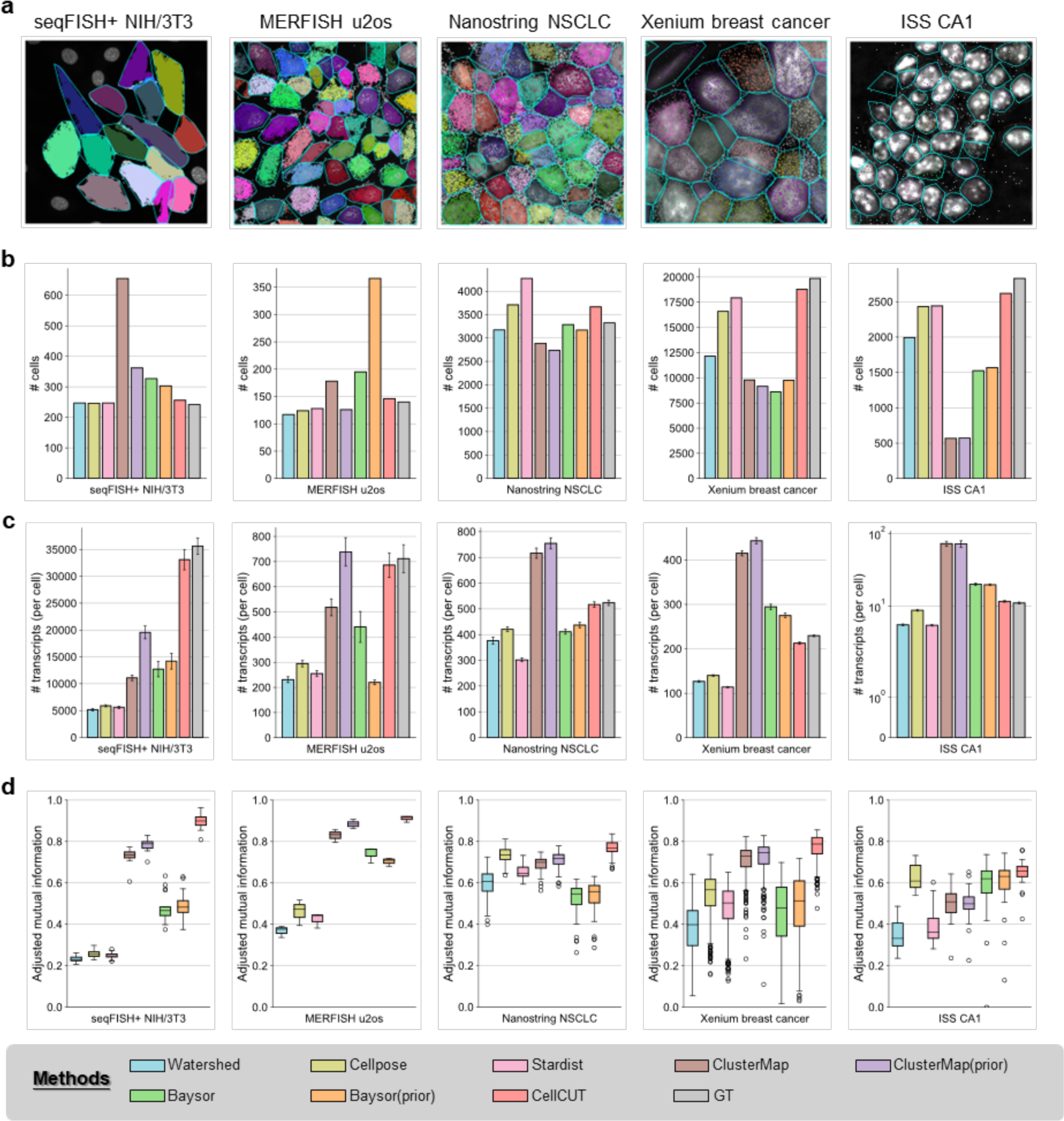
CellCUT provides precise cell segmentation performance. **a**, Example of segmentation results obtained by CellCUT in five datasets. The gray dots represent unassigned transcripts, different colors indicate different assignment results, and the cyan lines indicate the cell boundaries produced by CellCUT. The performances of CellCUT and eight competing methods in three evaluation benchmarks, which are: **b**, Cell counts, **c**, The average number of transcripts per cell, and **d**, The AMI metric, respectively. Error bars indicate the standard error of the average transcripts for each cell. Each AMI value is calculated in 512×512 pixels image in the NanoString NSCLC dataset and the Xenium breast cancer dataset, respectively. In the boxplot, the center line and the bounds of box refer to median, upper and lower quartiles, the whisker is equal to 1.5x interquartile range, and the dots are outliers.

We first evaluated CellCUT in the seqFISH+ NIH/3T3 dataset and the MERFISH u2os dataset. The seqFISH+ NIH/3T3 dataset included 17 fields of view (FoVs), and the MERFISH u2os dataset included 3 FoVs, each with a nucleus staining image of 2048×2048 pixels. In both datasets, most segmentation methods approximately counted the numbers of cells because the nuclei were clearly distinguishable, expect for Baysor and ClusterMap, which produced too many cells, as the individual cell was incorrectly segmented into multiple fragments (Supplementary Fig. S1). ClusterMap(prior) and Baysor(prior) slightly improved AMI metric over those of ClusterMap and Baysor in the seqFISH+ NIH/3T3 dataset, respectively, which suggested that incorporating the nucleus prior benefited the cell segmentation. However, the AMI of Baysor(prior) in the MERFISH u2os dataset was slightly lower than that of Baysor (0.76 versus 0.71), indicating that there is still room for improvement in cell segmentation models that explicitly model the affinities between nuclei and transcripts. The image-based methods (e.g., Watershed, Cellpose and Stardist) yielded overly conservative segmentation results, with fewer transcripts assigned to the cells, underestimated cell counts, and ignored the cell morphology, compared to the manual annotations (Fig. 2b-d).

We then applied CellCUT to NanoString NSCLC dataset and Xenium human breast cancer dataset, where cells were dense and irregular in shape. Accurately segmenting of individual cell with dense nuclei poses more challenging due to their close proximity, similar textures, and indistinct borders. Cellpose and Stardist generated the unstable results, due to image quality limitations (Supplementary Fig. S2-S3). In the Xenium breast cancer dataset, at least 9.7% of the cells were missed by deep learning models (Cellpose: 16,600, Stardist: 17,939, GT: 19,864), potentially due to low contrast intensity (Supplementary Fig. S3). The molecule-based methods performed unsatisfactorily cell profiles and miss a large number of cell (Supplementary Fig. S2-S3). Specifically, both Baysor and Baysor(prior) had average AMIs below 0.6 (Fig. 2d), indicating that cell boundaries were not accurately captured. The Kruskal-Wallis h-test was performed on multiple AMI metric and revealed significant differences between CellCUT and the other methods (P value < 2.7 × 10^-6^, n = 64 in the NanoString NSCLC dataset; P value < 2.3 × 10^-21^, n = 285 in the Xenium breast cancer dataset).

In contrast to the seqFISH+ NIH/3T3 dataset, the ISS CA1 dataset presented a distinct scenario, encompassing 96 genes with an average of approximately 10 transcripts per cell. Compared with other methods, CellCUT successfully identified 2,616 cells (Fig. 2b), closest matching the GT (2,828 cells). Stardist, the second-highest performer, identified 2,440 cells. The average AMI of CellCUT reached 0.65 (Fig. 2d), surpassing those of all image-based methods by at least 3.2% (Watershed: 0.35, Cellpose: 0.63, Stardist: 0.39). The molecule-based methods performed poorly with sparse transcripts, resulting in a severely insufficient number of cells (ClusterMap: 568, ClusterMap(prior): 572, Baysor: 1,524, Baysor(prior): 1,568). This underscored the generalizability of CellCUT, which provided stable segmentation results even with very dense or very sparse transcripts. Overall, the experimental results in Fig. 2 demonstrated that, by jointly leveraging nucleus staining images and spatial information acquired from transcripts, CellCUT clearly outperformed all compared methods from different platforms, cellular morphologies, and gene throughput, promising to recover cell counts and profiles that are much closer to the manually annotated cells.

### CellCUT is a robust model for undesirable data quality

Good data quality was crucial for cell segmentation performance, but real-world data often suffered from issues such as uneven image illumination, localized absence of transcripts, and blurred images. To demonstrate the effectiveness and robustness of the CellCUT, we conducted tests focusing on above three key aspects. First, we evaluated its performance in the STARmap mouse primary visual cortex brain region (VISp) dataset^4^ (Fig. 3a&3d, Supplementary Fig. S4-S5), comprising 1,020 genes with fewer than half a million transcripts. Qualitatively, it was clear from the nuclear staining image that there were some cells with low contrast and transcript enrichment that were not identified by Cellpose, and were successfully identified by CellCUT (Fig. 3a). Quantitatively, CellCUT identified 1,542 cells, higher than image-based method (Fig. 3d). Baysor and Baysor(prior) generated excessive cells without respecting the nucleus locations, as cells may be incorrectly divided into multiple fragments (Supplemental Fig. S4). Analysis of cell type proportions (Fig. 3g, Supplementary Fig. S5) revealed that Baysor mainly over-segmented Pvalb and L5 PT cells. These results highlighted the ability of CellCUT to effectively recover cell boundaries by combining nuclear segmentation and transcript information, overcoming limitations of insufficient segmentation in image-based methods and inaccurate boundaries in molecule-based methods.

**Fig. 3.**
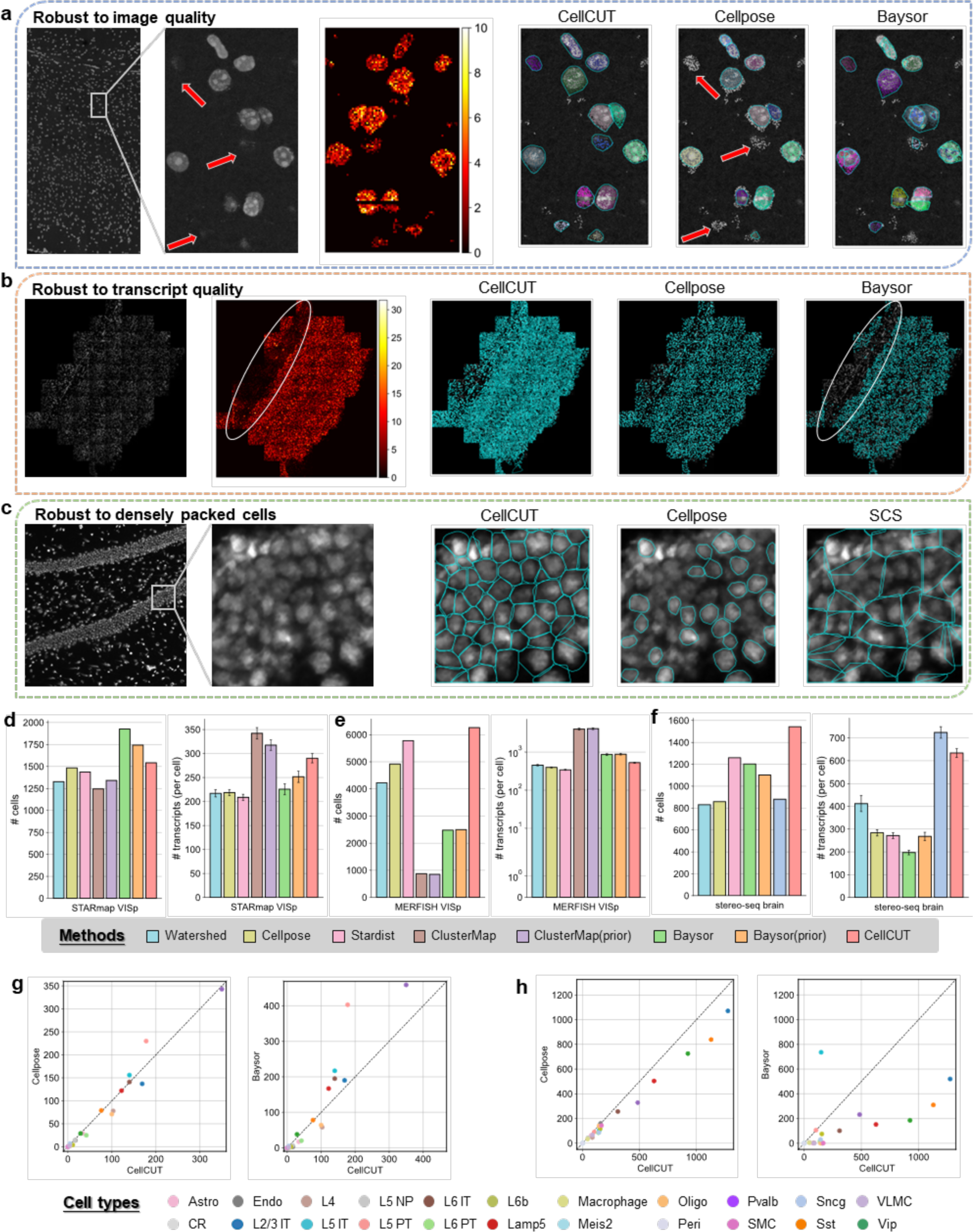
CellCUT is robust to undesired data quality. **a**, Example of cell segmentation results with low contrast image in the STARmap VISp dataset. The red arrows represent the cells that are not identified by Cellpose. **b**, Example of cell segmentation results with low transcript quality in the MERFISH VISp dataset. **c**, Example of segmentation results for densely packed cells in the Stereo-seq brain dataset. **d**, **e** and **f**, Quantitative comparisons of CellCUT to competitive methods in cells counts [left], and the average number of transcripts per cell [right] in the STARmap VISp dataset, in the MERFISH VISp dataset, and in the Stereo-seq brain dataset, respectively. **g** and **h**, Scatter plots showing the number of each cell type in the STARmap VISp dataset and the MERFISH VISp dataset. From left to right are scatter plot between Cellpose (y-axis) and CellCUT (x-axis), and scatter plot between Baysor (y-axis) and CellCUT (x-axis), respectively.

We then evaluated the performance of CellCUT in the MERFISH VISp dataset^24^ (Fig. 3b&3e, Supplementary Fig. S6-S7), comprising 268 genes with over three million transcripts, and decoding noise due to stack projection intensities. From the heat map of transcript counts, it was clear that there was a region of low transcript quality where molecule-based methods miss cellular structures (Fig. 3b, Supplementary Fig. S6). Quantitatively, 6,006 cells were identified by CellCUT (Fig. 3e), surpassing the second-highest performer, Stardist, by approximately 4.0% (5,777 cells), with 58.5% more transcripts per cell. Analysis of the proportions of cell types (Fig. 3h, Supplementary Fig. S7), revealed that CellCUT identified more L2/3 IT, Sst and Vip cells than other methods, while Baysor failed to identify many cell types, such as Oligo and VLMC.

Finally, we examined the segmentation performance on Stereo-seq^25^ (Fig. 3c&3f, Supplementary Fig. S8-S9), a sequencing-based ST technology that yields densely packed spots with a 0.5 µm spot-to-spot distance and an average of 3.3 unique molecular identifiers per spot, complemented by a single-stranded DNA staining image. Fig. 3c depicted a low-quality, densely packed tissue sample from the mouse hippocampal region. ClusterMap was excluded from the benchmark experiments due to its inapplicability to Stereo-seq, and SCS was included for comparison, a molecule-based cell segmentation method tailored for sequencing-based ST. The quantitative results (Fig. 3f) revealed that CellCUT identified a notably greater number of cells than other methods (Watershed: 831, Cellpose: 859, Stardist: 1,260, Baysor: 1,203, Baysor(prior): 1,102, SCS: 881, CellCUT: 1,543). Both Baysor and Baysor(prior) struggled to recover the appropriate cell size, resulting in limited transcripts per cell, primarily due to noisy spots (Supplementary Fig. S8). SCS was sensitive to initial nucleus segmentation, and tended to merge multiple cells into one in the dentate gyrus region, underestimating cell counts (Supplementary Fig. S9). CellCUT accurately recovered cellular structures at the dentate gyrus region (Supplementary Fig. S9), with 22.5% more cells and 133.9% more transcripts per cell than the second-highest performer. Overall, CellCUT jointly optimized image segmentation and transcript information, reduced the impact of undesirable data quality, and therefore outperforming other competitive methods.

### CellCUT reveals the fine spatial structures to multiple ST platforms

We then illustrated that the accurate cell segmentation of CellCUT can reveal the fine spatial structures to ST platforms in the NanoString NSCLC dataset and the Xenium human breast cancer dataset. By counting four comprehensive evaluation metrics, we observed that CellCUT identified more cells and assigned more total transcripts in terms of overall characteristics (Fig. 4a&4c). The proportion of assigned transcripts in the Cellpose and Baysor was less than 60% and 80% of the CellCUT results, respectively. In terms of cell-level metrics, the number of transcripts and genes assigned to individual cells of CellCUT was also superior to other methods. Although the average number of transcripts was slightly higher in cells segmented by Baysor, the gene diversity per cell less than CellCUT. We then compared the correlation between cell type signatures segmented by CellCUT in the CosMx and scRNA-seq platforms for NanoString NSCLC dataset (Fig. 4b), and in the Xenium and Chromium platforms for human breast cancer dataset (Fig. 4d), and observed that the average expression between spatial and sequencing data in both datasets was highly correlated. Additional correlation heatmaps from other segmentation results can be found in Supplementary Fig. S12 for comparison.

**Fig. 4.**
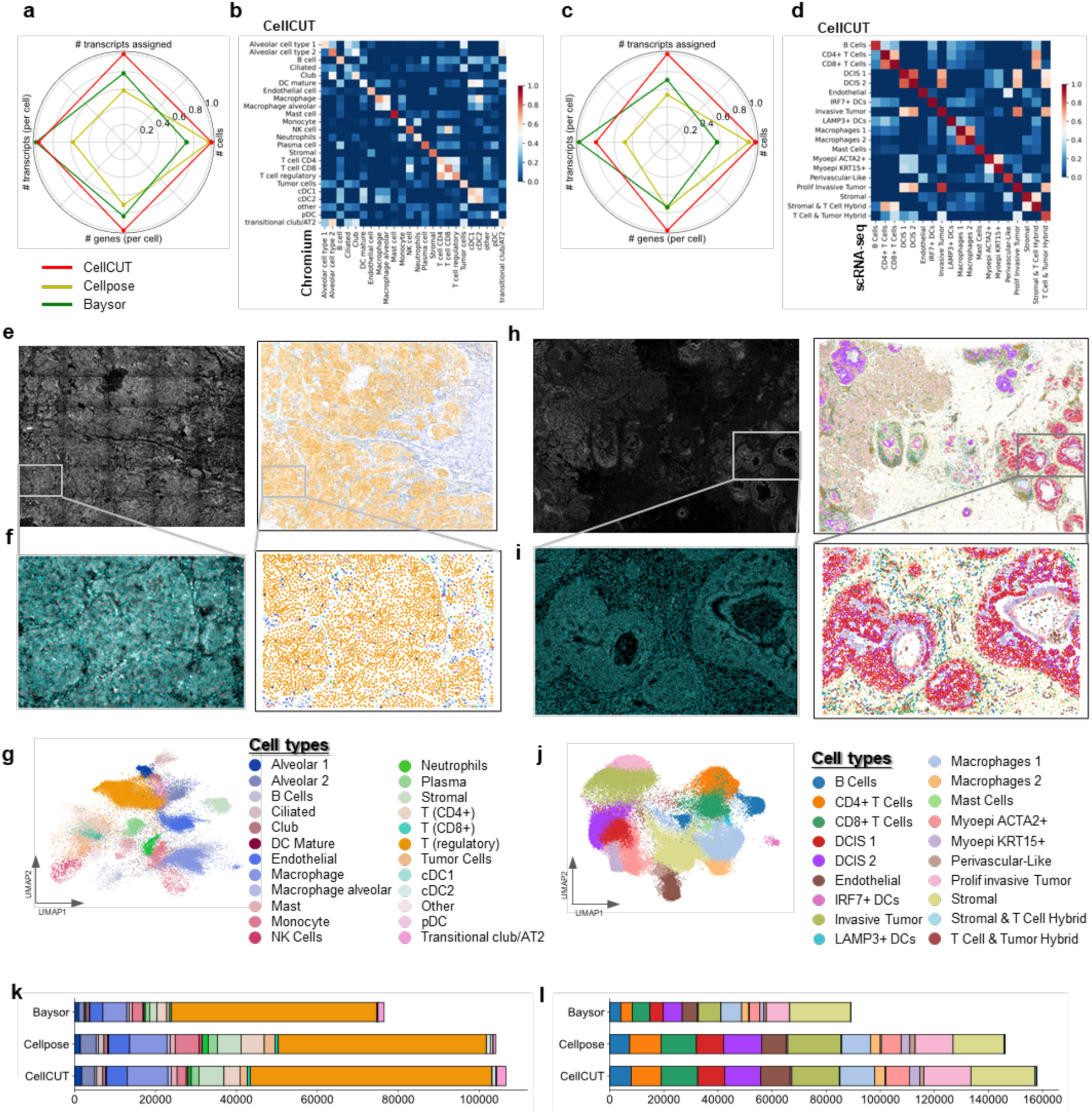
CellCUT reveals the fine spatial structures to multiple ST platforms. **a**, Radar plot shows the comparison of four evaluation metrics in the NanoString NSCLC dataset. Each color represents a different method. **b**, Correlation heatmap of average expression between segmented cells from CellCUT (y-axis) and expression from Chromium data (x-axis) in the NanoString NSCLC dataset. **c**, Radar plot shows the comparison of four evaluation metrics in the Xenium breast cancer dataset. Each color represents a different method. **d**, Correlation heatmap of average expression between segmented cells from CellCUT (y-axis) and expression from scRNA-seq data (x-axis) in the Xenium breast cancer dataset. **e**, Nucleus staining image and the spatial distributions of all identified cells by CellCUT in the NanoString NSCLC dataset, where each dot represents the segmented cells colored by the cell type annotation. **f**, Zoomed-in views of cell boundaries and cell type annotations. **g**, UMAP of segmented cells colored based on cell type annotation. **h**, Nucleus staining image and the spatial distributions of all identified cells by CellCUT in the Xenium breast cancer dataset. **i**, Zoomed-in views of cell boundaries and cell type annotations. **j**, UMAP of segmented cells colored based on cell type annotation. **k** and **l**, The number of each cell type identified by different segmentation methods in the NanoString NSCLC dataset and the Xenium breast cancer dataset, respectively.

In the NanoString NSCLC dataset, CellCUT, Cellpose, and Baysor all successfully identified two major cell types including macrophage and tumor cells, and showed distinct spatial distributions (Fig. 4e-4g, Supplementary Fig. S10). Unlike CellCUT, Cellpose and Baysor missed a number of tumor cells characterized by densely packed nuclei and unclear borders (Supplementary Fig. S10). In the Xenium human breast cancer dataset, CellCUT, Cellpose, and Baysor all identified circular duct-like structures, with DCIS cells densely packed together and surrounded by myoepithelial cells (Fig. 4h-4j, Supplementary Fig. S11). Differently, CellCUT discovered more myoepithelial cells characterized by irregular nuclei morphology (Supplementary Fig. S11). Moreover, CellCUT identified more endothelial and stromal cells compared to Cellpose, which were characterized by elongated shapes, low contrast intensity, and therefore not easily recovered by image-based methods. Baysor severely underestimated the number of cells for almost all cell types due to the excessive boundaries recovered by Baysor (Supplementary Fig. S11). Overall, CellCUT effectively recovered cell boundaries by integrating nucleus segmentation and transcript information to overcome the insufficient segmentation of image-based methods and the inaccurate boundaries of molecule-based methods, thereby revealing more detailed spatial structures.

### CellCUT integrates a human-in-the-loop workflow to generalize to customized dataset

The pretrained CNN model demonstrated its generalizability across diverse nucleus staining images. A challenge remained in tailoring cell segmentation for customized data with substantial stylistic differences from publicly available datasets. To address this, we introduced a human-in-the-loop pipeline for quickly obtaining high-quality pixel embeddings (Fig. 5a). With new images, CellCUT(pretrained) was first run to generate initial nucleus segmentation results. An annotator then corrected the segmentation errors and delineated missing or inaccurate regions. Following this, a new finetuned model was trained on these manually annotated images, and applied to subsequent images in the dataset. The annotator iteratively refined the newly nucleus segmentation, retraining the pixel embeddings model with the annotated images. This process continued until the annotator was satisfied with the accuracy or the manual corrections converged to zero. Leveraging the robust pretrained model, this loop typically required only a few iteration (typically less than 3, empirically) to achieve satisfactory results.

**Fig. 5.**
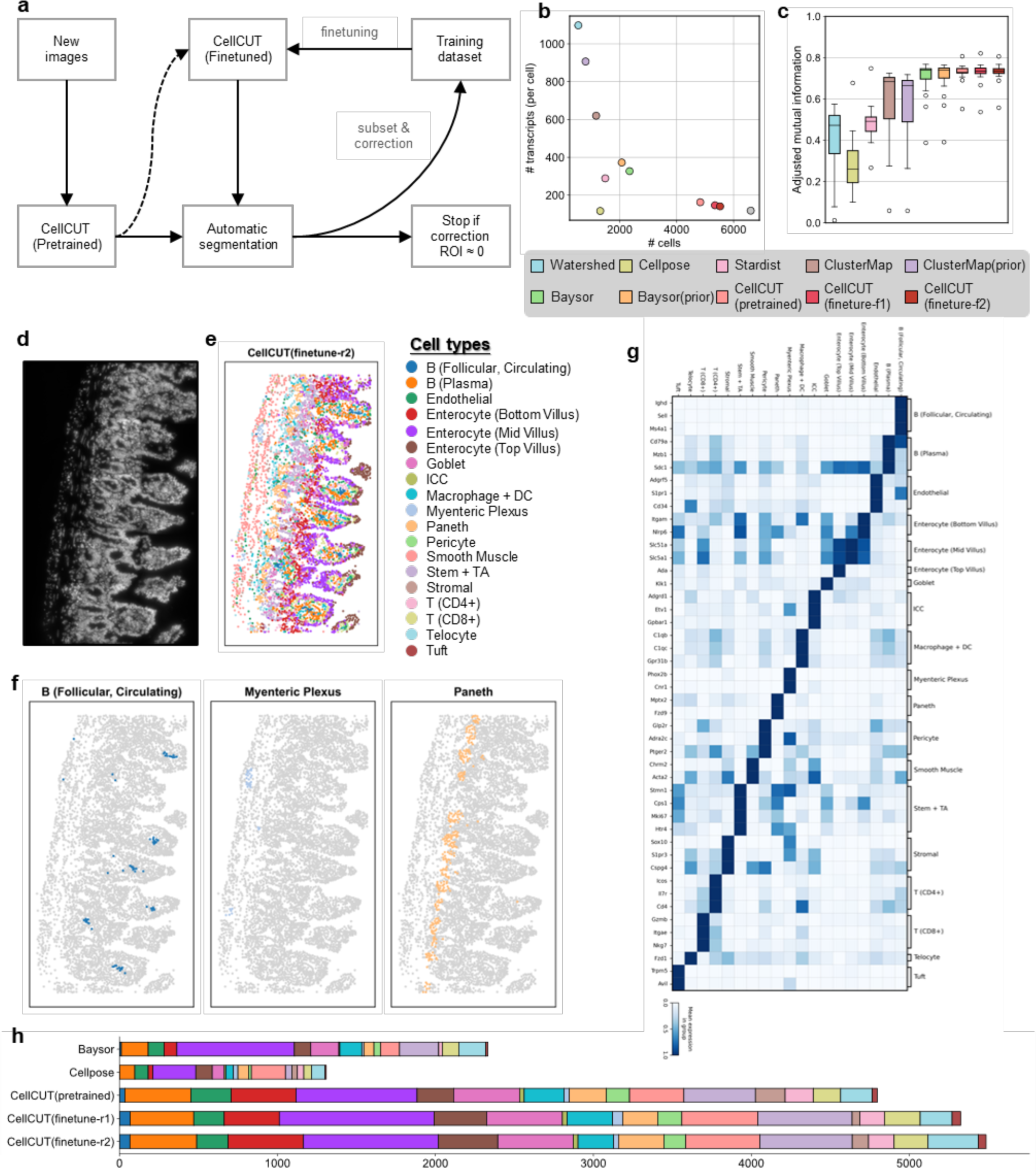
CellCUT integrates a human-in-the-loop workflow to enhance its generalizability to customized datasets. **a**, Schematic of the human-in-the-loop procedure. **b**, Scatter plot showing the average transcripts per cell against the cell counts. Each dot represents a different method. **c**, The AMI metric obtained across all benchmark studies, each AMI value is calculated in 512×512 pixels image. In the boxplot, the center line and the bounds of box refer to median, upper and lower quartiles, the whisker is equal to 1.5x interquartile range, and the points are outliers. **d,** Nucleus staining image of MERFISH mouse ileum dataset. **e**, Spatial distributions of all identified cells by CellCUT (finetune-r2) colored based on cell type annotation. **f**, Three representative rare cell types. **g**, Expressions of marker genes in each of the identified cell clusters. The size of the dots represents the fraction of cells with at least one count of the indicated gene. The color of the dots represents the average expression of each gene across all cell types. **h**, The number of each cell type identified by different segmentation methods.

We applied this workflow in the MERFISH mouse ileum data^20^ (Fig. 5d, Supplementary Fig. S13-S15), a dataset in which cell segmentation is challenging due to uneven illumination and densely packed cells. Two rounds of manual correction were performed, each time annotating two 1024×1024 pixels nucleus segmentations, fine-tuning the pretrained model, and running CellCUT for cell segmentation. First, we compared CellCUT(pretrained) with other benchmark methods to evaluate the pretrained model (Fig. 5b-5c). Overall, we found that CellCUT outperformed other methods, both in terms of the number of cells and the captured transcripts per cell. To evaluate the finetuned models, we compared cell counts, average transcripts per cell, and AMIs among CellCUT(finetune-r1), CellCUT(finetune-r2) and CellCUT(pretrained). Quantitatively, the number of identified cells increased (pretrained: 4,828, finetune r1: 5,345, finetune r2: 5,524, GT: 6,601), the number of average transcripts per cell slightly decreased (pretrained: 164, finetune-r1: 146, finetune-r2: 142, GT: 118), and the average AMI metric remained roughly invariant (approximately 0.73), implying that the finetuned model improved segmentation performance as rounds increased (Fig. 5b-5c). We next conducted the single cell analysis of three representative segmentation methods to clarify the types of segmented cells (Fig. 5h, Supplementary Fig. S13). Cellpose segmented only a small number of cells and failed to identify B (follicular, circulating) cells, struggling to recover detailed spatial structure. Similarly, Baysor identified only 42.5% of the cells identified by CellCUT(finetune-r2) and failed to identify stromal cells (Supplementary Fig. S13). Compared to image-based methods and molecule-based methods, CellCUT(finetune-r2) reproduced the diverse cell types expected in the mouse ileum (Fig. 5e, Supplementary Fig. S15), detected more cells, and recovered more rare cell types, such as tuft cells, myenteric plexus, B (Follicular, Circulating) cells, and ICC cells (Fig. 5f, Supplementary Fig. S13). CellCUT(finetune-r1) and CellCUT(finetune-r2) yielded similar results (Supplementary Fig. S14), with the number of individual cell types identified mostly higher than that of CellCUT(pretrained). There result demonstrated that the finetuned CellCUT was able to recover more rare cell types, contributing to cell type annotation and quantitative modeling of tissue architectures.

### CellCUT enables subcellular analysis of spatial transcriptomics

High-resolution spatial transcriptomics provided an opportunity to explore the molecular heterogeneity within individual cells. CellCUT naturally divided the segmentation results of each cell into nuclear and cytoplasmic regions, identifying subcellular structures, providing valuable insights into RNA localization and composition at the subcellular level (Fig. 6a, Methods). We performed t-tests to identify genes localized differently in the two regions, and found many genes were significantly enriched in the two identified regions (Fig. 6b-e). To visualize the spatial distribution of genes, Supplementary Fig. S16 show selected visualization results of tagging the top three genes with the most significant differences for each category. In the SeqFISH+ NIH/3T3 dataset (Fig. 6b), the marked genes were derived from the original paper^9^, and in the MERFISH u2os dataset (Fig. 6c) the marked genes were from Bento^26^, both of which revealing consistency and agreement between the two results. Notably, the KPNB1 gene was predominantly localized in the nucleus, and is crucial for nucleocytoplasmic transport of macromolecules. The PSAT1 gene encoded the phosphoserine aminotransferase 1 enzyme involved in the serine biosynthesis pathway, and is primarily localized in the cytoplasm of cells. The TLN1 gene played a role in cell adhesion and migration within the cytoplasm. The CKAP5 gene, which contributed to the dynamic regulation of microtubule stability, was expressed in the cytoplasm.

**Fig. 6.**
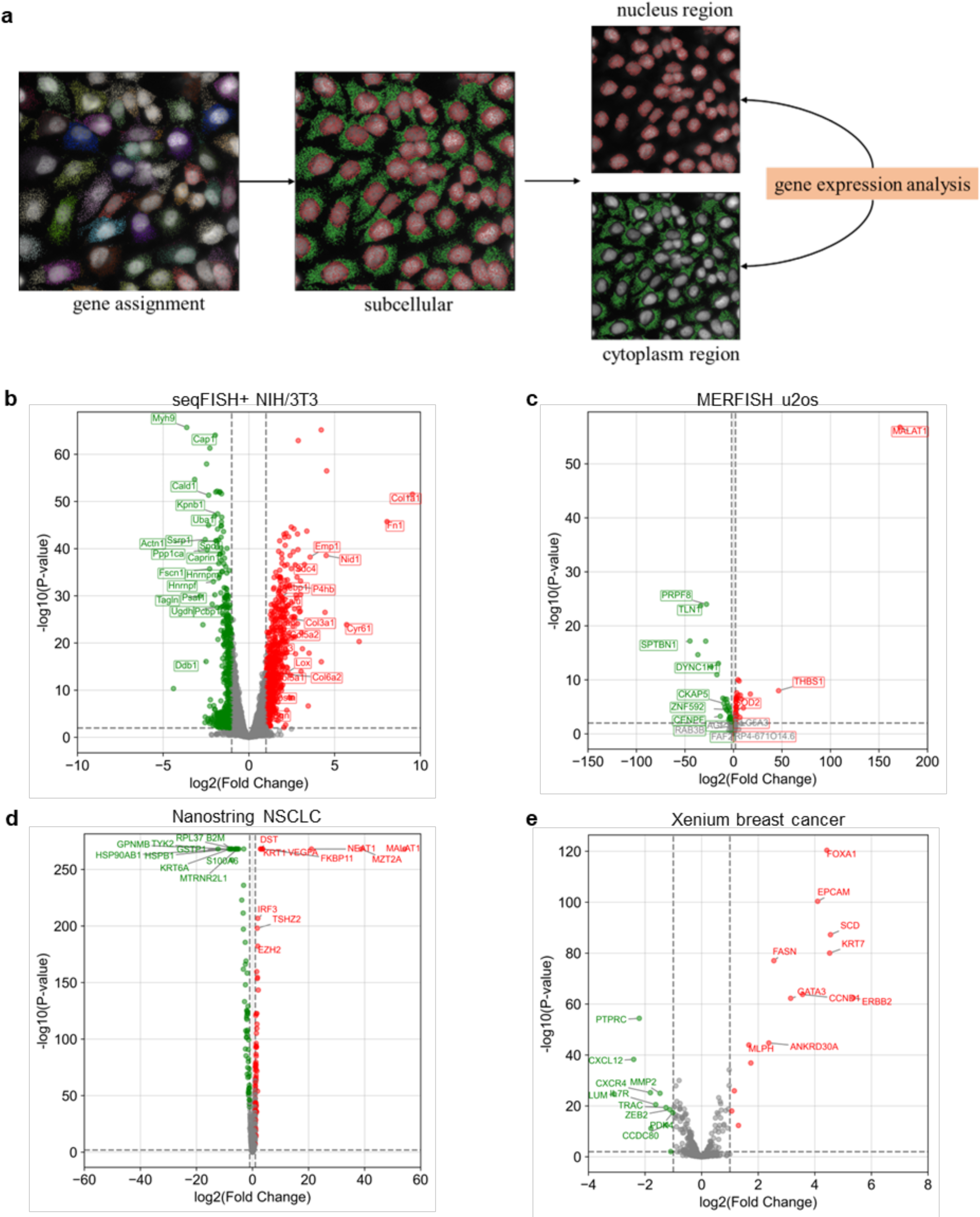
CellCUT enables subcellular analyses of spatial transcriptomics. **a**, Schematic diagram of subcellular analyses by dividing transcripts into nuclear and cytoplasmic regions. b, Volcano plot that shows quantitative changes by the expression levels of the genes between the nucleus and cytoplasm of CellCUT segmented cells for the seqFISH+ NIH/3T3 dataset. Genes with P values < 0.01 and fold changes greater than 1 are identified from each group (t-tests, one-sided). c, Volcano plot for the MERFISH u2os dataset. Genes with P values < 0.01 and fold changes greater than 2 are identified from each group (t-tests, one-sided). d, Volcano plot for the NanoString NSCLC dataset. Genes with P values < 0.01 and fold changes greater than 2 are identified from each group (t-tests, one-sided). The top 10 genes with the smallest P values are identified from each group (t-tests, one-sided). e, Volcano plot for the Xenium breast cancer dataset. Genes with P values < 0.01 and fold changes greater than 2 are identified from each group (t-tests, one-sided). The top 20 genes with the smallest P values are identified from each group (t-tests, one-sided). RNA genes enriched in the nucleus or cytoplasm are colored. The red dots indicate genes enriched in the nucleus, the green dots indicate genes enriched in the cytoplasm, and the grey dots indicate genes with no significant differences.

In the NanoString NSCLC dataset (Fig. 6d) and the Xenium breast cancer dataset (Fig. 6e), we tagged the genes with the most significant differences, which were in agreement with the prior knowledge in gene subcellular locations. For example, the long noncoding RNA genes Neat1 and MALAT1, known nuclear transcripts that formed core components of organelles in the nucleus. The TYK2 gene, involved in cytokine signaling, primarily localized to the cytoplasm. The PTPRC gene predominantly localized on the plasma membrane of immune cells, regulating immune cell activation and signaling. Overall, these results demonstrated that CellCUT efficiently identified subcellular structures, facilitating subcellular analyses of high-resolution spatial transcriptomics data. Importantly, subcellular analysis with CellCUT requires no additional computational burden, demonstrating the utility of CellCUT in parsing spatial transcriptomic data.

## Discussion

Spatial transcriptomics, a rapidly advancing technology, has deepened our understanding of cellular organization by revealing the intricate spatial distributions of the different cell types within tissues. This detailed knowledge, particularly in the brain, sheds light on the physiological functions of distinct cell populations and their intricate signaling interactions. However, cell segmentation, the crucial step for parsing these spatial data, remains a major challenge, hindering the full potential of ST technology.

Current cell segmentation methods in ST fall short, relying solely on staining images or molecules, leading to inaccurate segmentation results or missed cells. We develop CellCUT, a novel framework that combines both data sources, to address these limitations. CellCUT leverages a pretrained model to capture spatial cues from nuclei, and then refines transcript affinities via supervised learning. Finally, the superpixels and transcript within a constrained graph are jointly optimized to produce cell segmentation results.

We performed a comprehensive evaluation on multiple datasets derived from a diverse dataset of ST platforms. CellCUT detected more cells, assigned more transcripts to individual cells, and yielded more accurate cell borders than eight widely used segmentation methods. In addition, further analysis of the cell type annotation results indicated that CellCUT was able to recover a greater number of rare cell types, benefiting other downstream applications. The analysis of the spatial distribution of RNAs revealed that many RNAs were enriched in different subcellular regions, which is consistent with the prior knowledges. These findings demonstrated the ability of CellCUT to facilitate high-resolution spatial transcriptomics at both the cellular and subcellular levels.

The ability of CellCUT to accurately and robustly recover cells from ST data comes from the fact that CellCUT leverages both nuclear information from staining images and gene expression profiles from transcripts. Segmentation methods that rely only on staining images usually produce more conservative segmentation results, and struggle with ambiguous areas and overlapping cells. However, segmentation methods relying only on transcripts tend to segment similarly distributed transcripts into a single cell, ignoring the morphological information of the cell, and are sensitive to the density of the given transcripts. Unlike these methods, CellCUT effectively fuses these two types of information through a joint optimization framework, which is able to recover more reasonable cell boundaries as well as overcome the limitations of single cues.

CellCUT facilitates model retraining through a human-in-the-loop process unlocks a unique advantage in situations where the pretrained model falls short in terms of nucleus segmentation. This iterative approach allows for tailored model refinement, significantly enhancing the generalizability of the model. Even in the face of diverse cell morphologies, CellCUT can be finetuned by correcting unsatisfactory nuclei, leading to improved cell segmentation overall.

Although CellCUT outperform other competitive segmentation algorithms, several potential ways can be used to further improve its cell segmentation performance. First, augmenting the number of initial transcript features can enhance the affinity prediction between transcripts. This includes considering shape features of cells, topological features of transcripts, or incorporating additional prior information like cell type annotations. Second, deep learning can be used to automatically extract discriminative transcript features in the future, thereby optimizing the efficiency of the algorithm. In addition, the hyperparameters for the graph partitioning model are presently determined empirically. Although the model exhibits robustness to these hyperparameters, automating their determination could further reduce manual intervention time. Finally, CellCUT is a model-agnostic computational framework, so both the CNN and GAT can be replaced with more advanced network structures, like the transformer network.

In summary, we presented a framework called CellCUT for cell segmentation in ST data. Application to a range of datasets involving diverse tissues and ST protocols showed CellCUT outperformed state-of-the-art algorithms, achieving superior performance without the need for manual annotation. CellCUT is effective in densely packed cellular regions, detecting more cells and yielding more accurate cell borders, thereby contributing to the recovery of rare cell types and fine-grained spatial structures. We believe that CellCUT can serve as a very useful tool in high-resolution ST data, facilitating detailed downstream cellular and subcellular analyses, and generating novel biological insights.

## Methods

### CNN model for nucleus prediction

#### Model architecture for image

The CNN model is a deep neural network with a U-net^27^ style and incorporates residual blocks^28^. The network takes a single-channel nucleus staining image as its input and yields two outputs: the affinity of a pixel to nearest neighbor pixels^29^ (2d represent x- and y- respectively), and a high-dimensional representation of pixels (24d in our experiment).

#### Loss function

Since the network has multi-task outputs, the loss function consists of two terms. For the first term, we use a weighted sum of the binary cross-entropy loss and the Dice loss to measure the difference between the predicted affinity map and the ground truth affinity map. For the second term, we adopt the discriminative loss^30^ to push or pull embeddings of each pixel.

#### Nucleus training datasets

Several publicly available nucleus datasets are collected to train the pretrained CNN model. These datasets consist of 2,511 images.

### GNN model for transcript embeddings prediction

#### Initial features extraction

The initial features for each transcript serve as the input of the GNN model, comprising two components, named neighborhood gene composition (NGC) and neighborhood embedding composition (NEC).

#### Model architecture for transcript

We adopt the graph attention network^33^ (GAT), a well-known GNN that introduced the attention mechanism, to update the initial transcript features.

### Graph partitioning model

Assuming that nuclei within the same cells and their transcripts are similar, this allows the construction of a joint optimization model to recover cell. The model construction is divided into three steps.

#### Construct the superpixels graph

Upon completing the nucleus prediction, we can construct a superpixels graph through the affinity map and embeddings map to represent the nucleus segmentation.

#### Construct the transcripts graph

Upon completing the transcript embeddings prediction, we can construct a graph to represent the similarities of transcripts.

#### Construct the whole graph

Upon obtaining the superpixels graph and transcripts graph, we construct a whole graph to perform cell segmentation jointly.

#### Optimization model

The optimization problem is equivalent to partitioning *G_C_*, with the goal of dividing the nuclei and transcripts of the same cell into the same classes. This problem can be solved efficiently by a heuristic greedy solver^38^ with polynomial time complexity. Nodes belonging to the same component in the partition result are assigned one cell. Finally, the cell borders can be estimated by calculating the convex hull of the nucleus segmentation and transcript assignment for visualization. This procedure is performed separately for each component.

### Comparison of Methods

Several representative cell segmentation methods were used to compared, including image-based approaches (Watershed, Cellpose, Stardist) and molecule-based approaches (Baysor, ClusterMap, SCS).

#### Watershed

Watershed is utilized for segmentation based on nuclei staining images with three steps. First, the nuclei masks are first obtained by OTSU thresholding. Subsequently, the masks are processed using the distance transform algorithm to generate markers by peak detections. Finally, the masks and markers are used as input for the watershed algorithm, to produce the final cell segmentation.

#### Cellpose

Cellpose is a popular generalist cell segmentation method based on the U-net. It provides several pretrained models for different image styles. In our benchmark study, we use the pretrained model (including ‘cyto’, ‘nuclei’, ‘cyto2’, ‘CP’, ‘CPx’) with the best performance and nuclei staining images for different datasets. Cell diameter is estimation automatically for different datasets.

#### Stardist

Stardist is a widely-used segmentation method based on the Mask R-CNN with shape priors. It provides three pertained models for fluorescent images. Similarly, we use the pretrained model with the best performance and nuclei staining images for different datasets for comparison.

#### Baysor

Baysor is a molecule-based method based on the MRF for ST data. It allows clustering of molecular with and without prior cell segmentation. Therefore, in our benchmark study, we evaluate the performance of Baysor both with and without prior segmentation. We choose the default configuration and adjust the parameter ‘scale’ according to the cell size in each dataset.

#### ClusterMap

ClusterMap is a molecule-based method for ST data. Its clusters molecules based on the physical location and gene identity of molecules by density peak clustering. Similarly, this method can incorporate staining images as an auxiliary input. In our benchmark study, we evaluate the performance of ClusterMap both with and without nucleus prior. We choose the default configuration and adjusted the parameter ‘xy_radius’ according to the cell size in each dataset.

#### SCS

SCS is original designed for sequencing-based ST data. It assigns spots to cells by a transformer network to learn the position of each spot relative to the center of its cell. In our benchmark study, we utilize SCS with the default configuration in the Stereo-seq dataset.

### Evaluation metrics

We evaluated the different algorithms in three aspects: cell counts, average number of transcripts per cell, and AMI compared with ground truth. In the case of image-based segmentation methods, we converted their instance results to transcript assignments for a fair comparison. Therefore, cell counts were determined by the number of unique cluster IDs in the assignment results, representing the number of segmented cells. The average number of transcripts per cell was calculated as the count of transcripts within each cluster, indicating an estimate of the average size per cell. AMI, a common metric for unsupervised clustering evaluation, was used to measure the similarity between the assignment results and the ground truth cluster IDs, with higher values indicating better segmentation accuracy. For fair and stable performance, cells with transcript counts less than a constant were excluded from the evaluation metric calculation.

### Single cell analysis

Single-cell analysis was performed by using Scanpy^39^. All methods segmented cells adopted the same following filter criteria for a robust performance: transcript counts of >20, and the number of genes expressed being ≥10. All genes expressed in at least 10 cells were kept for subsequent analysis. For MERFISH mouse ileum data, fifty principal components were then extracted, a neighbor graph was constructed by considering the 15 nearest neighbors for cells, a UMAP embedding was calculated, the Leiden algorithm was used for clustering, and subclustering was performed on some of the clusters to produce the final clusters. The marker genes were chosen following the original paper^20^. For MERFISH VISp dataset, STARmap VISp dataset, Stereo-seq brain dataset, NanoString NSCLC dataset and Xenium breast cancer dataset, we transferred cell type labels from scRNA-seq data to the segmented cells. In each dataset, we took the intersection of genes from the scRNA-seq data and the spatial data, and then used the ingest data integration function from the Scanpy package to perform the label transferring. For plot correlation heatmap, we calculated the Pearson correlation of the average expression for each cell type between the segmented cells and publicly available single-cell datasets.

### Subcellular analysis

Cell segmentation results produced by CellCUT can be divided into two groups: a nucleus group and a cytoplasm group. The nucleus group was identified by transcripts overlay in nucleus segmentation results, while the cytoplasm group is the remaining transcripts. To conduct subcellular analysis, transcripts within each group were summarized to form gene expression profiles, which were then aggregated by region into a gene matrix. We used t-tests to identify genes with different RNA localization between the two groups, and highlighted genes in each group that had a P value < 0.01 and a fold change greater than 1 (for the seqFISH+ NIH/3T3 dataset and the Xenium breast cancer dataset) or 2 (for the MERFISH u2os dataset and the NanoString NSCLC dataset). Genes enriched in the nucleus are indicated in red, and genes enriched in the cytoplasm are indicated in green.

### Data availability

All data used in evaluating the developed methods have been previously published, including:

1. seqFISH+ NIH/3T3 cells, 10,000 genes.
2. MERFISH u2os cells, 241 genes, in this dataset, the decoded genes were obtained by PIPEFISH^40^.
3. NanoString CosMx SMI human non-small-cell lung cancer, 980 genes.
4. 10X Genomics Xenium human breast cancer, 541 genes.
5. STARmap mouse VISp, 1,020 genes.
6. MERFISH mouse VISp, 268 genes.
7. ISS mouse CA1 region, 95 genes.
8. Stereo-seq mouse brain, 26,177 genes.
9. MERFISH mouse ileum, 241 genes.

### Code availability

Watershed implemented by skimage and opencv, open-source Python packages Cellpose (v.2.1.1), Stardist (v.0.8.3), Baysor (v.0.6.2), ClusterMap (v.0.0.0) and SCS (v.0.0.0) were applied for comparison with CellCUT.

---

**Supplementary Fig. S1.**
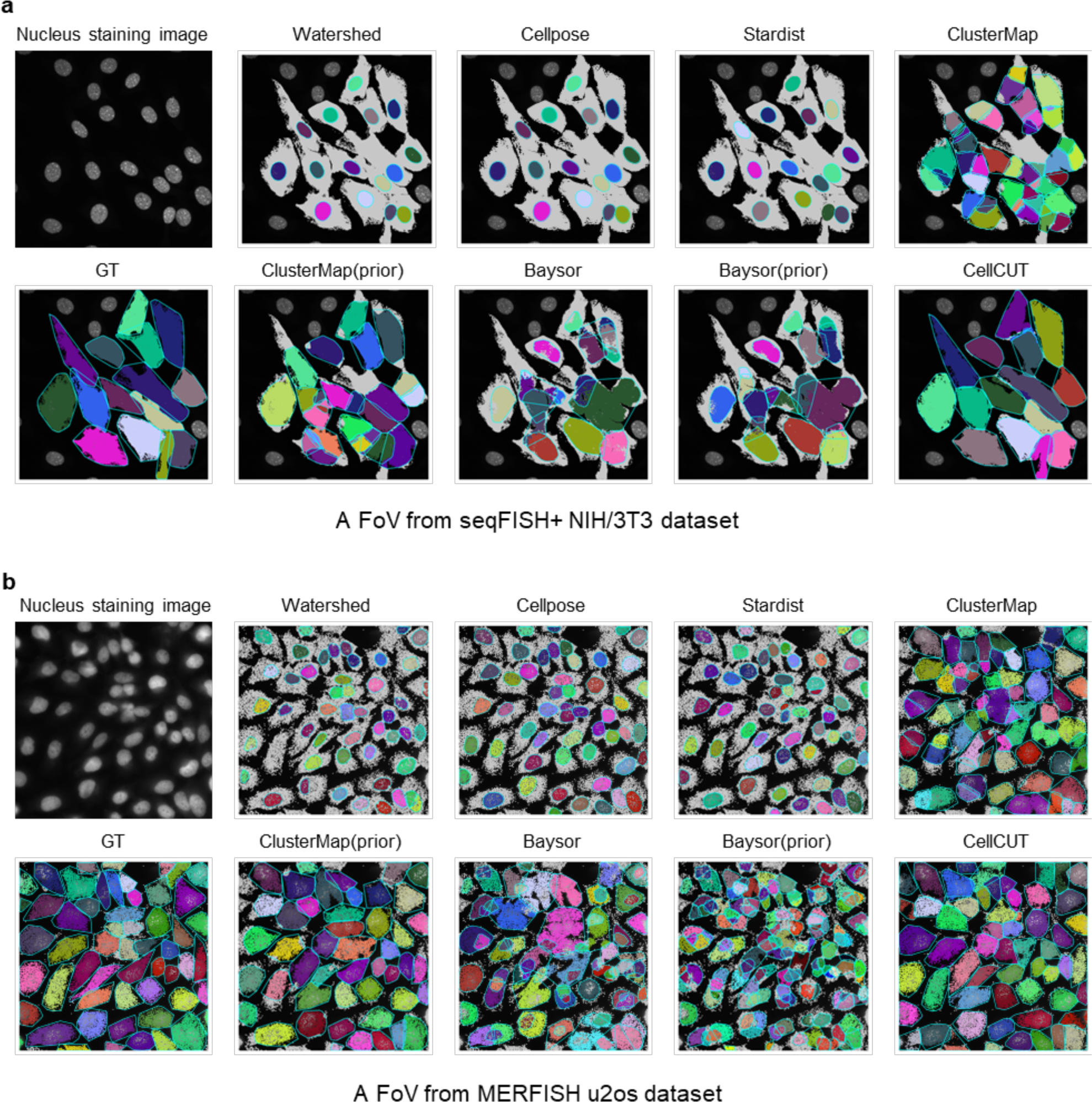
Visual comparisons of cell segmentation and transcript assignment results for **a**, FoV 5 in the SeqFISH+ NIH/3T3 dataset, and **b**, FoV 3 in the MERFISH u2os dataset are illustrated, the original nucleus staining image has 2048×2048 pixels. The cyan lines delineate the cell boundaries, the gray dots represent unassigned transcripts, and the dots with different colors indicate the different transcript assignment results.

**Supplementary Fig. S2.**
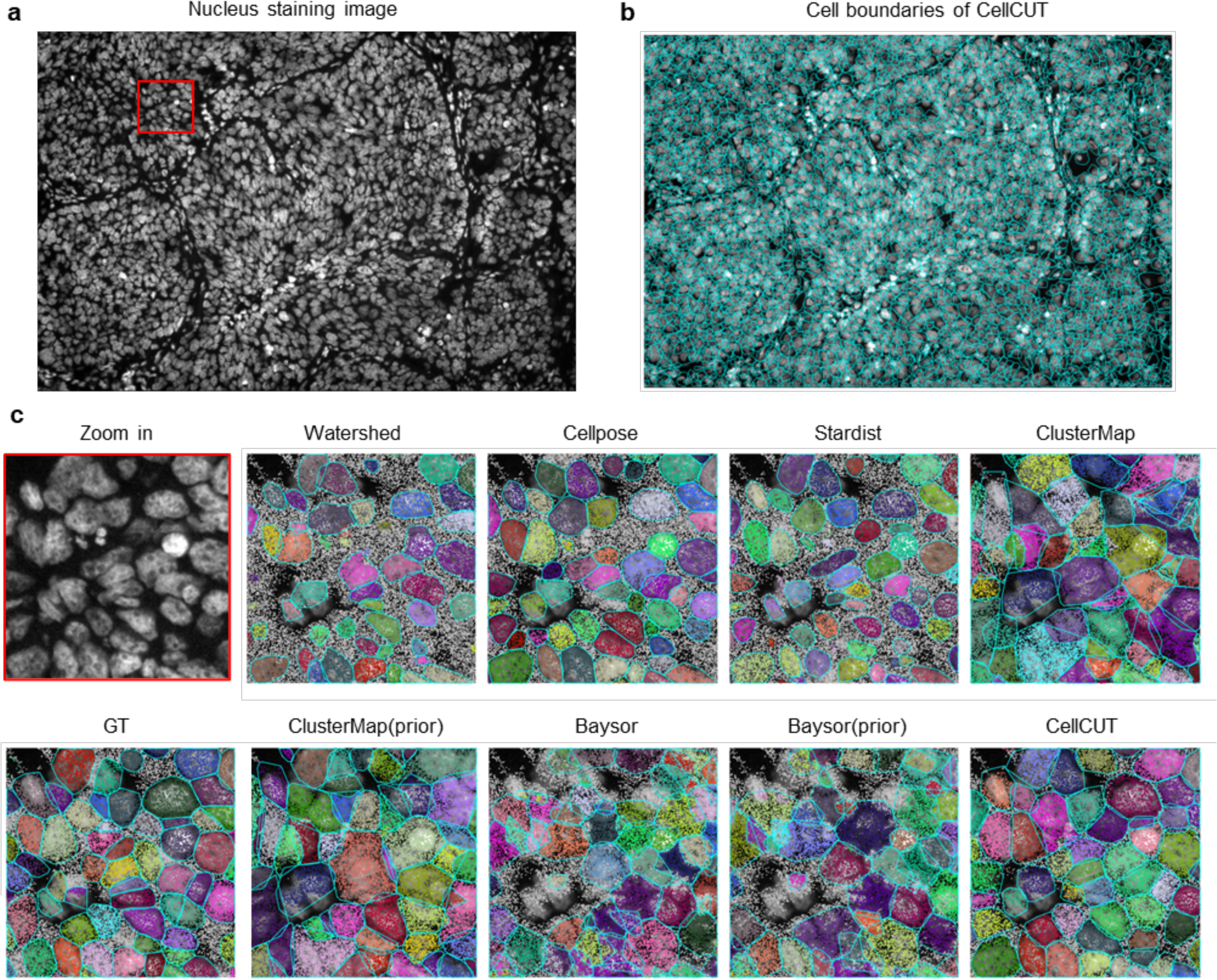
Visualizations of cell segmentation and transcript assignment produced by different segmentation methods in the NanoString NSCLC dataset. **a**, The original nucleus staining image has 5472×3678 pixels. **b**, Visualization of the cell boundaries produced by CellCUT. **c**, Zoomed-in views of a 256×256 pixels zone in the original image, the corresponding ground truth, and the visual comparisons of the different methods. The cyan lines delineate the cell boundaries, the gray dots represent unassigned transcripts, and the dots with different colors indicate the different transcript assignment results for each method.

**Supplementary Fig. S3.**
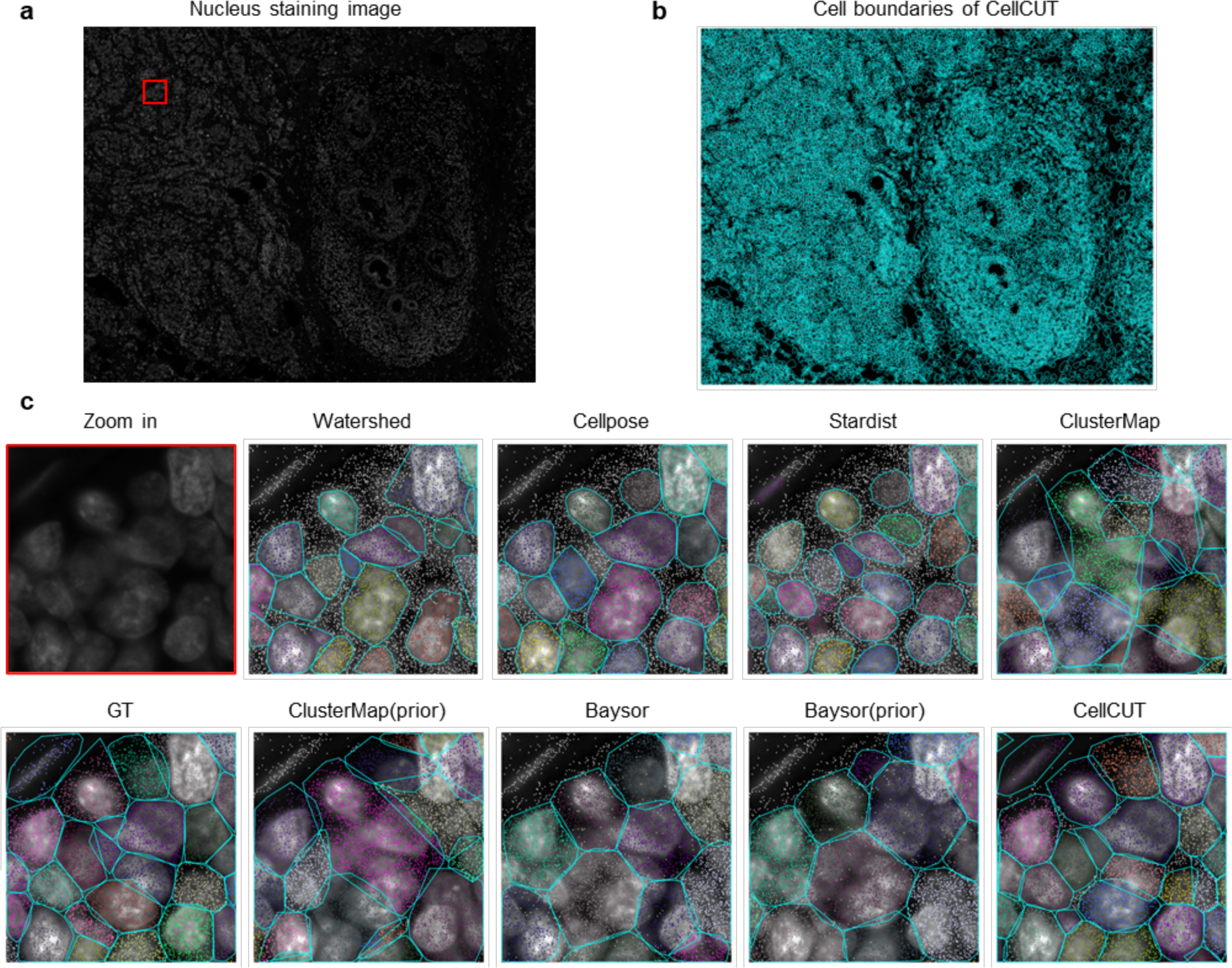
Visual comparisons of cell segmentation and transcript assignment results in the Xenium breast cancer dataset. **a**, A ROI of 9500×7500 cropped from the original nucleus staining image. **b**, Visualization of the cell boundaries produced by CellCUT. **c**, Zoomed-in views of a 128×128 pixels zone in the original image, the corresponding ground truth, and the visual comparisons of the different methods. The cyan lines delineate the cell boundaries, the gray dots represent unassigned transcripts, and the dots with different colors indicate the different transcript assignment results for each method.

**Supplementary Fig. S4.**
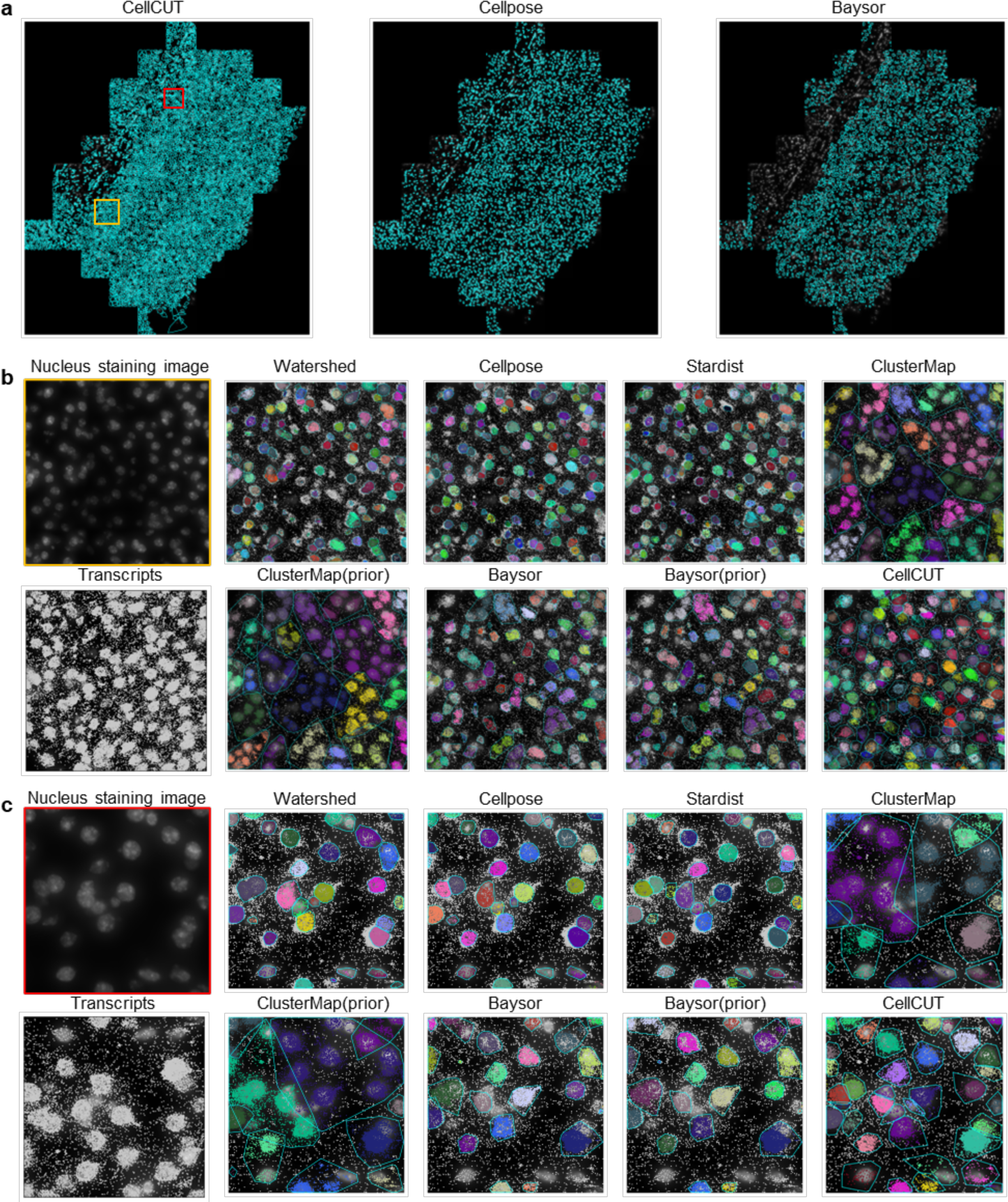
Visual comparisons of cell segmentation and transcript assignment results in the MERFISH VISp dataset. a, Visualization of the cell boundaries produced by CellCUT, Cellpose, and Baysor, respectively. **b** and **c**, Visual comparisons across CellCUT and other methods in the zoomed-in zone in a. The yellow and red boxes indicate the zoomed-in views, the cyan lines indicate cell boundaries, the gray dots represent unassigned transcripts, and the dots with different colors indicate the different transcript assignment results for each method.

**Supplementary Fig. S5.**
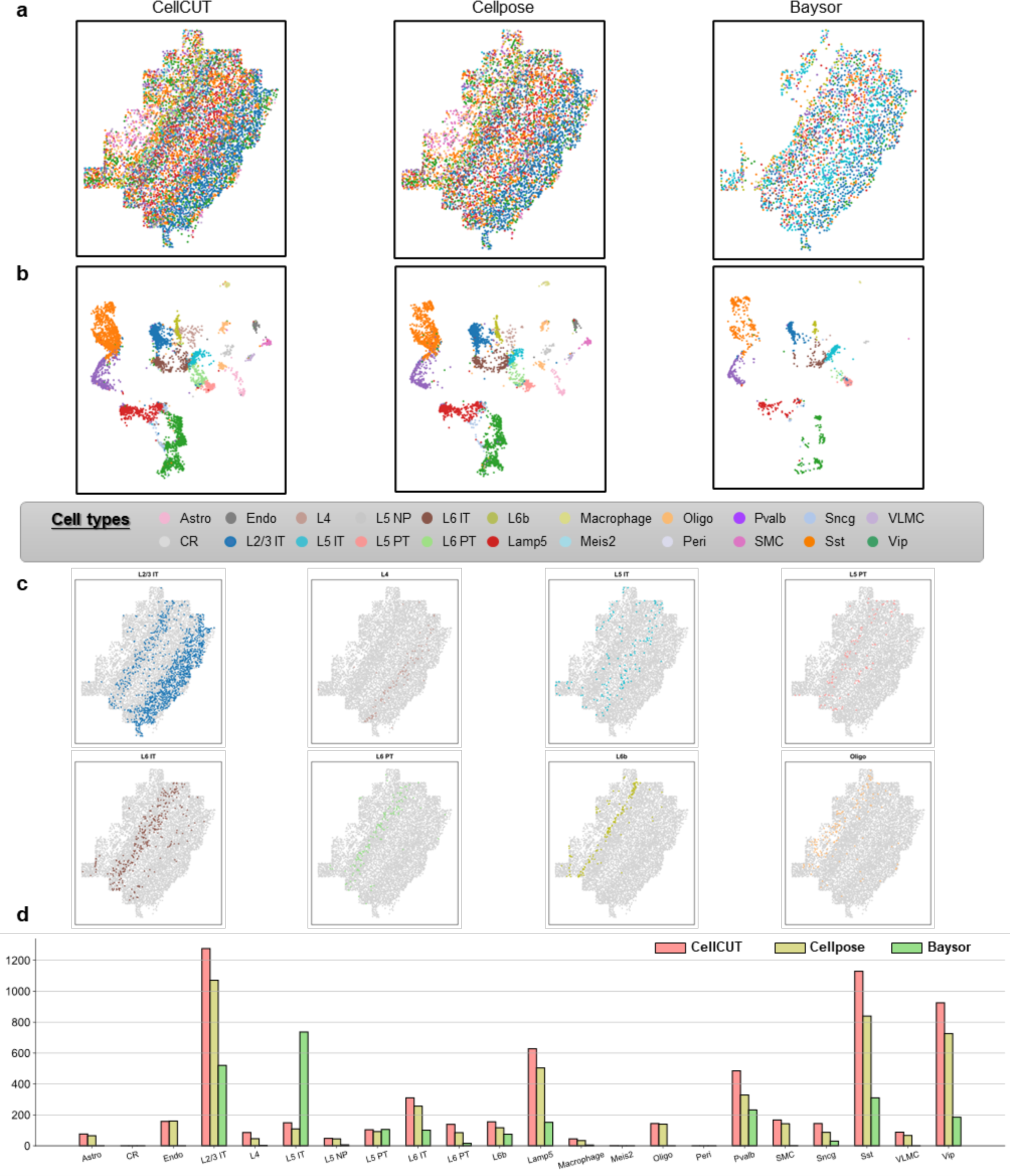
Comparisons of spatial structures in the MERFISH VISp dataset recovered from different methods. **a**, Spatial distributions of all identified cell. **b**, UMAP of segmented cells colored based on cell type annotation. From left to right, results produced by CellCUT, Cellpose, and Baysor, respectively. c, Spatial distributions of several cell types identified by CellCUT. The gray dots represent the location of all cells and the colored dots represent the location of the indicated cell type. **e**, The number of each cell type identified by each of the segmentation methods in a-d.

**Supplementary Fig. S6.**
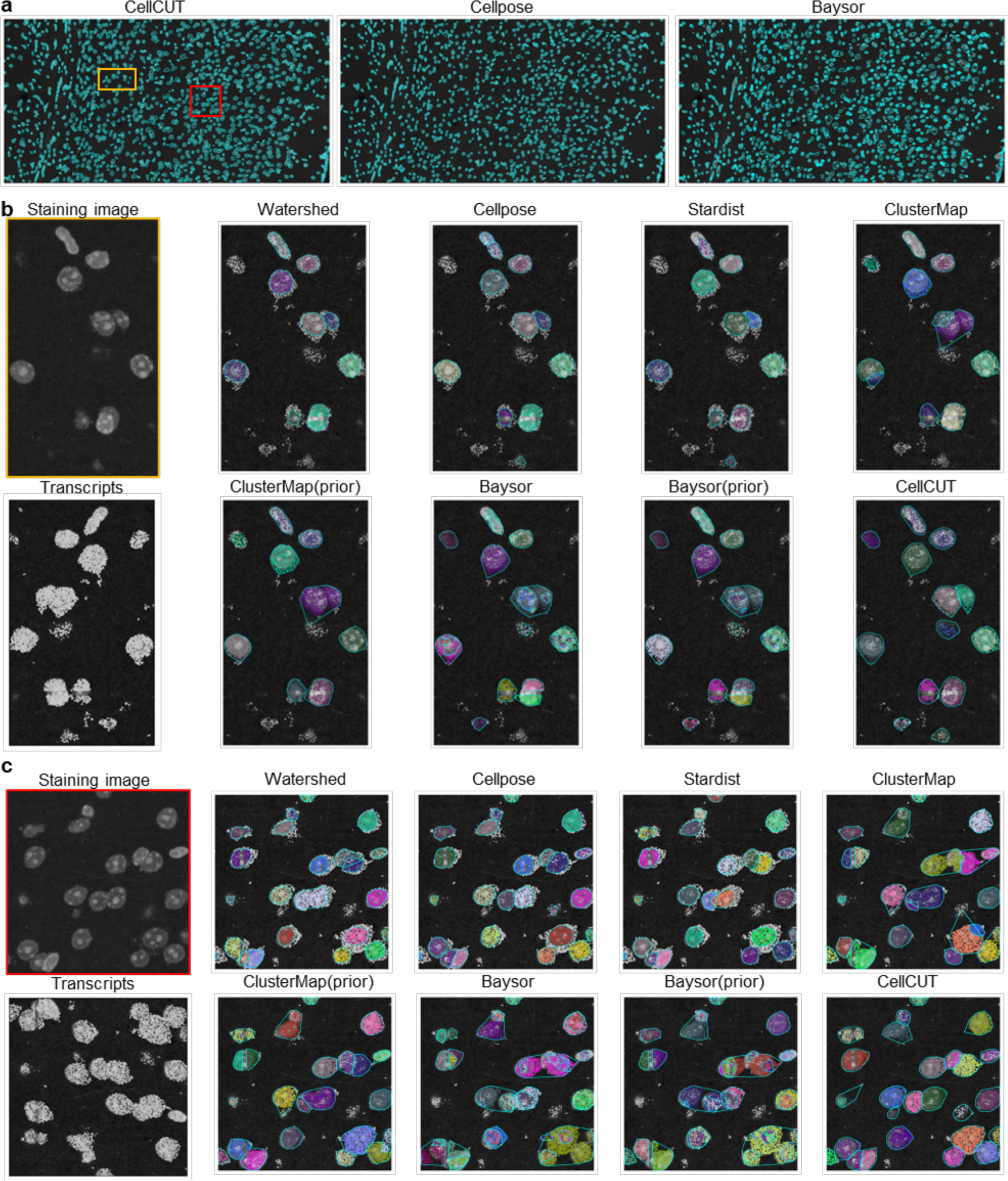
Visual comparisons of cell segmentation and transcript assignment results in the STARmap VISp dataset. a, Visualization of the cell boundaries produced by CellCUT, Cellpose, and Baysor, respectively. **b** and **c**, Visual comparisons across CellCUT and other methods in the zoomed-in zone in a. The yellow and red boxes indicate the zoomed-in views, the cyan lines indicate cell boundaries, the gray dots represent unassigned transcripts, and the dots with different colors indicate the different transcript assignment results for each method.

**Supplementary Fig. S7.**
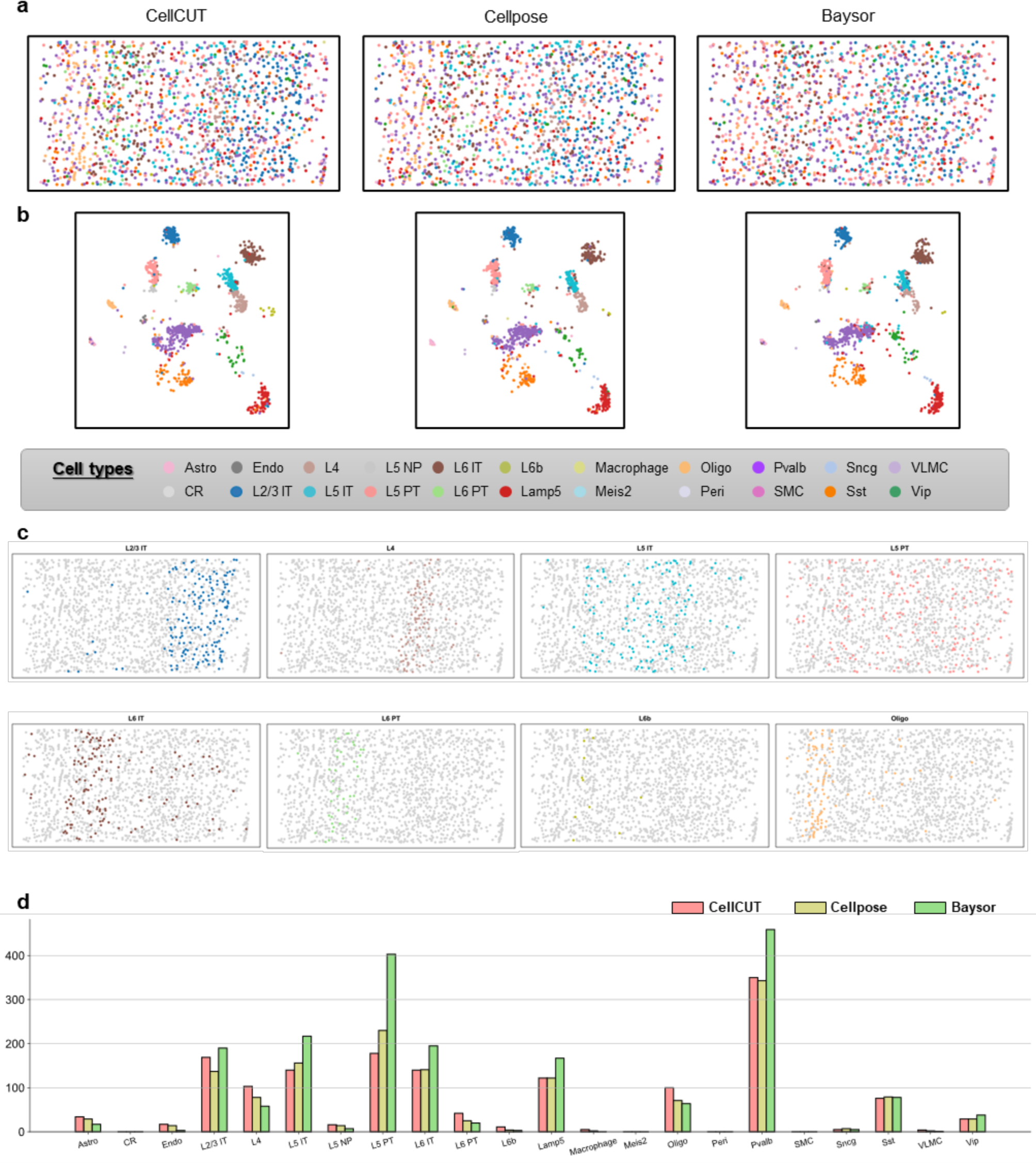
Comparisons of spatial structures in the STARmap VISp dataset recovered from different methods. **a**, Spatial distributions of all identified cell. **b**, UMAP of segmented cells colored based on cell type annotation. From left to right, results produced by CellCUT, Cellpose, and Baysor, respectively. c, Spatial distributions of several cell types identified by CellCUT. The gray dots represent the location of all cells and the colored dots represent the location of the indicated cell type. **e**, The number of each cell type identified by each of the segmentation methods in a-d.

**Supplementary Fig. S8.**
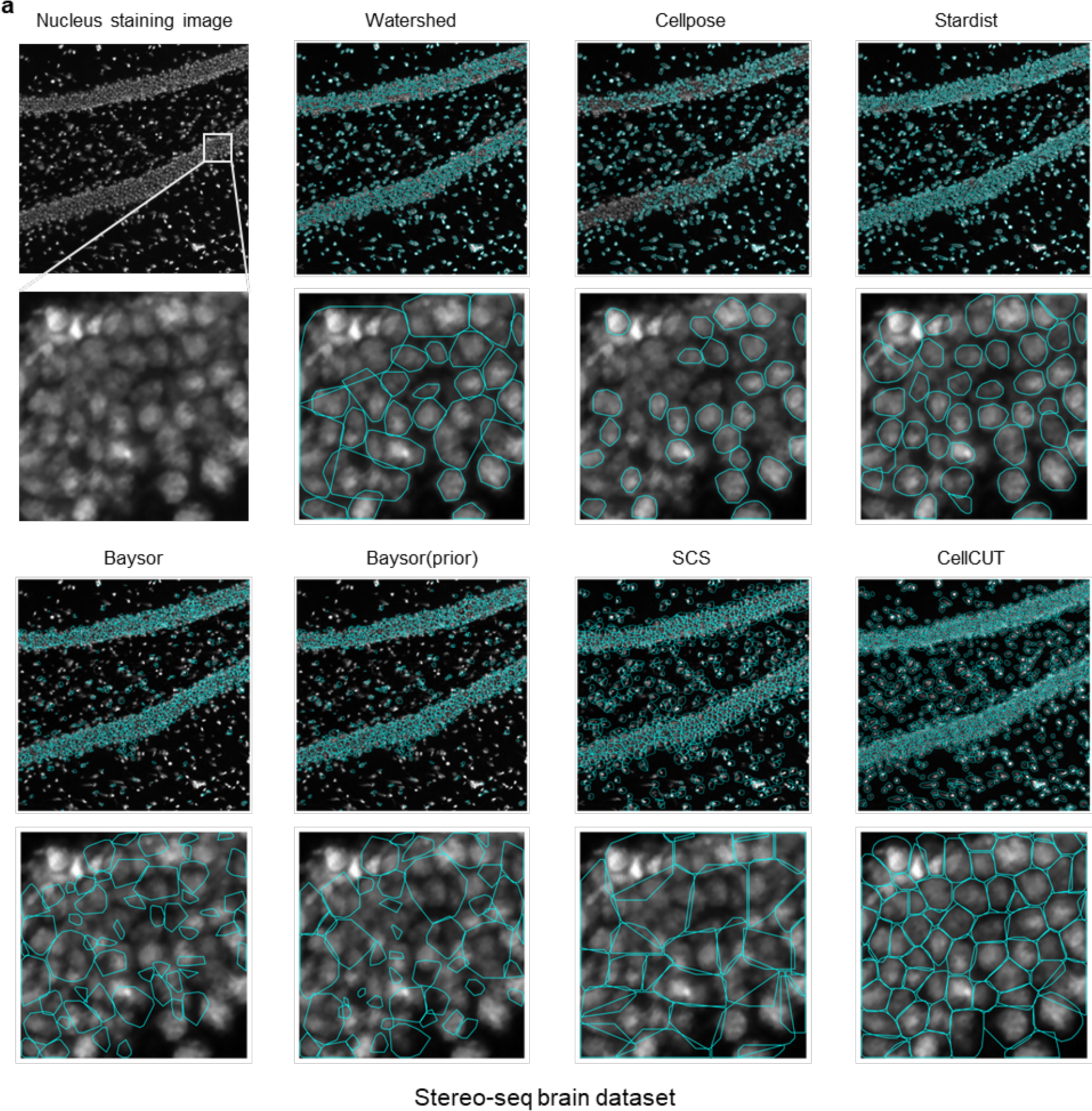
Visualization of cell segmentation results obtained by different segmentation methods in the Stereo-seq brain dataset. The original nucleus staining image has 1200×1200 pixels, the zoomed-in views have 128×128 pixels. The cyan lines indicate the cell boundaries for each method.

**Supplementary Fig. S9.**
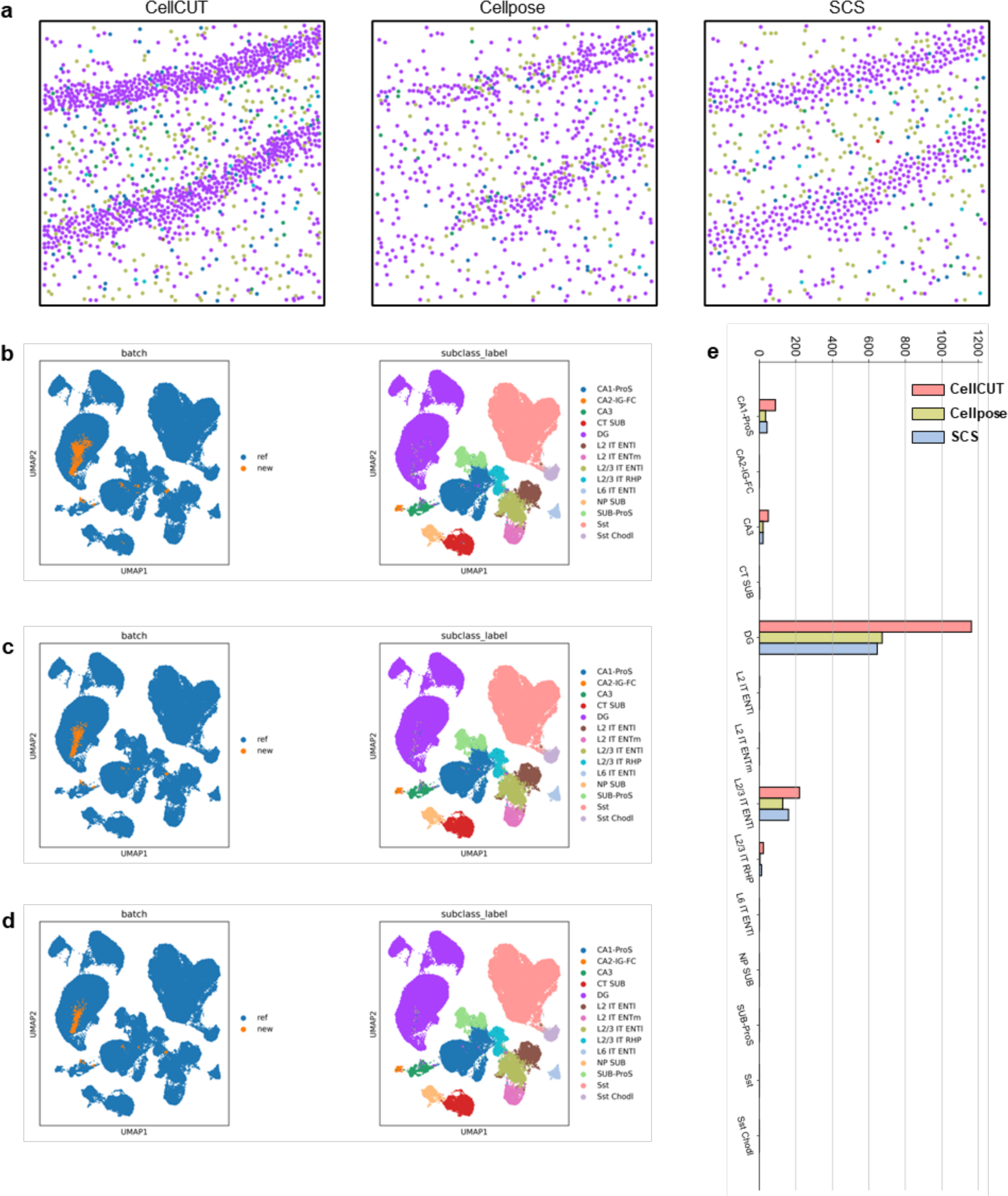
Comparisons of spatial structures in the Stereo-seq brain dataset. **a**, Spatial distributions of all identified cell recovered from CellCUT, Cellpose, and SCS, respectively. **b**, **c**, and **d** are UMAP of scRNA-seq and segmented cells from CellCUT, Cellpose, and SCS, respectively. **e**, The number of each cell type identified by each of the segmentation methods in a-d.

**Supplementary Fig. S10.**
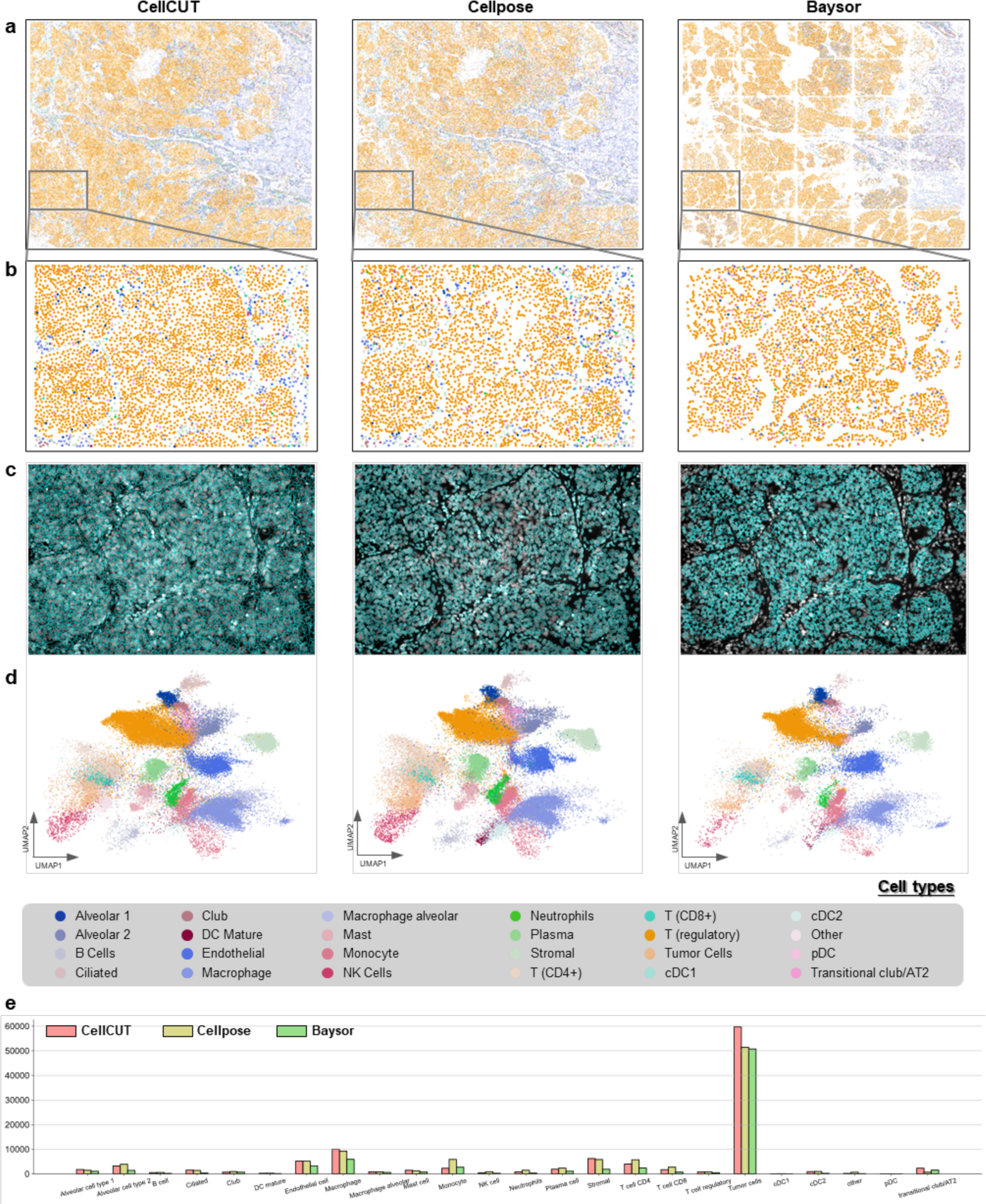
Comparisons of spatial structures in the NanoString NSCLC dataset recovered from different methods. **a**, **b**, **c**, and **d** are the cell type annotations, zoomed-in views of cell type annotations, cell boundaries, and UMAP of segmented cells colored based on cell type annotation, respectively. From left to right, results produced by CellCUT, Cellpose, and Baysor, respectively. **e**, The number of each cell type identified by each of the segmentation methods in a-d.

**Supplementary Fig. S11.**
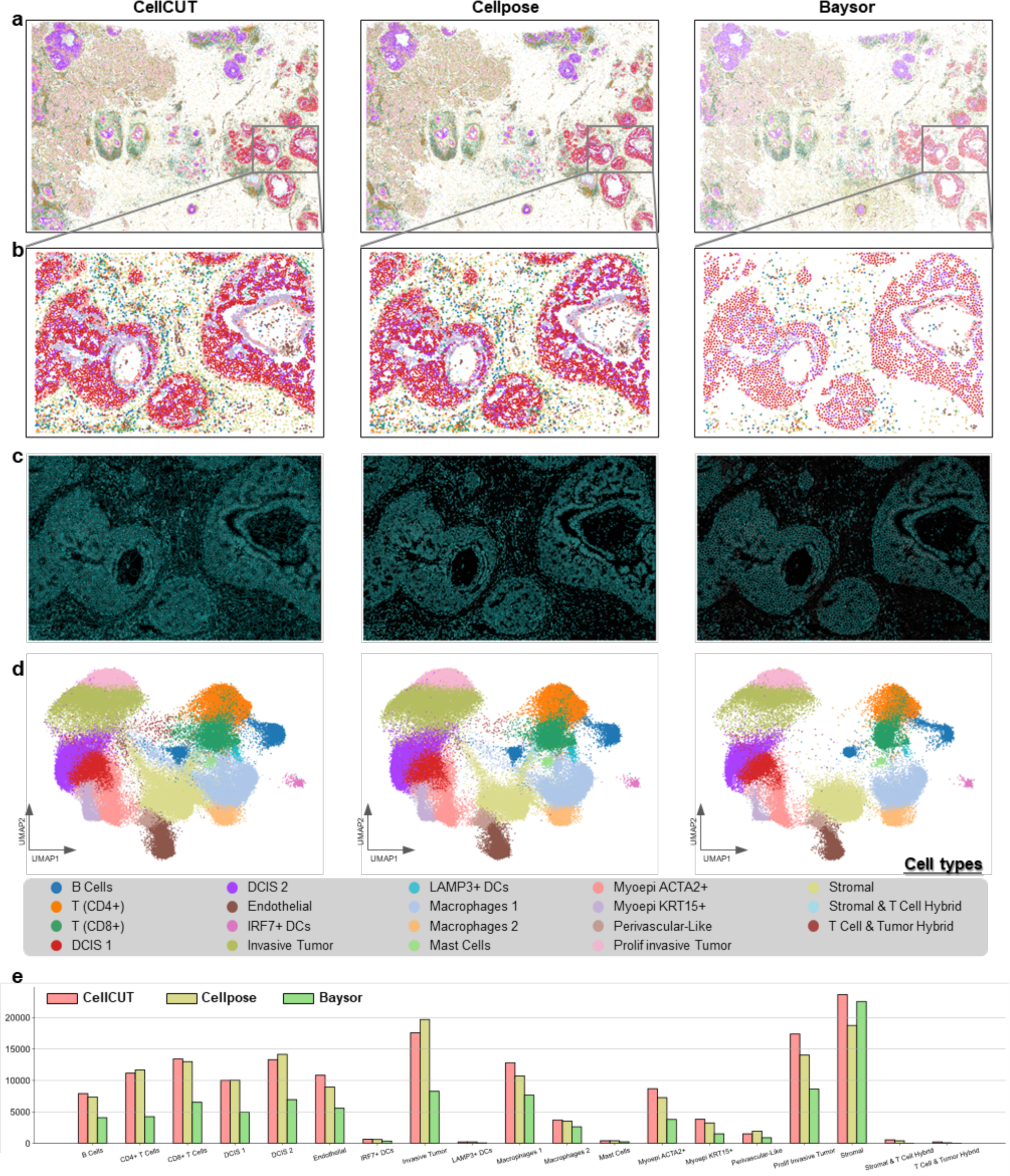
Comparisons of spatial structures in the Xenium breast cancer dataset recovered from different methods. **a**, **b**, **c**, and **d** are the cell type annotations, zoomed-in views of cell type annotations, cell boundaries, and UMAP of segmented cells colored based on cell type annotation, respectively. From left to right, results produced by CellCUT, Cellpose, and Baysor, respectively. **e**, The number of each cell type identified by each of the segmentation methods in a-d.

**Supplementary Fig. S12.**
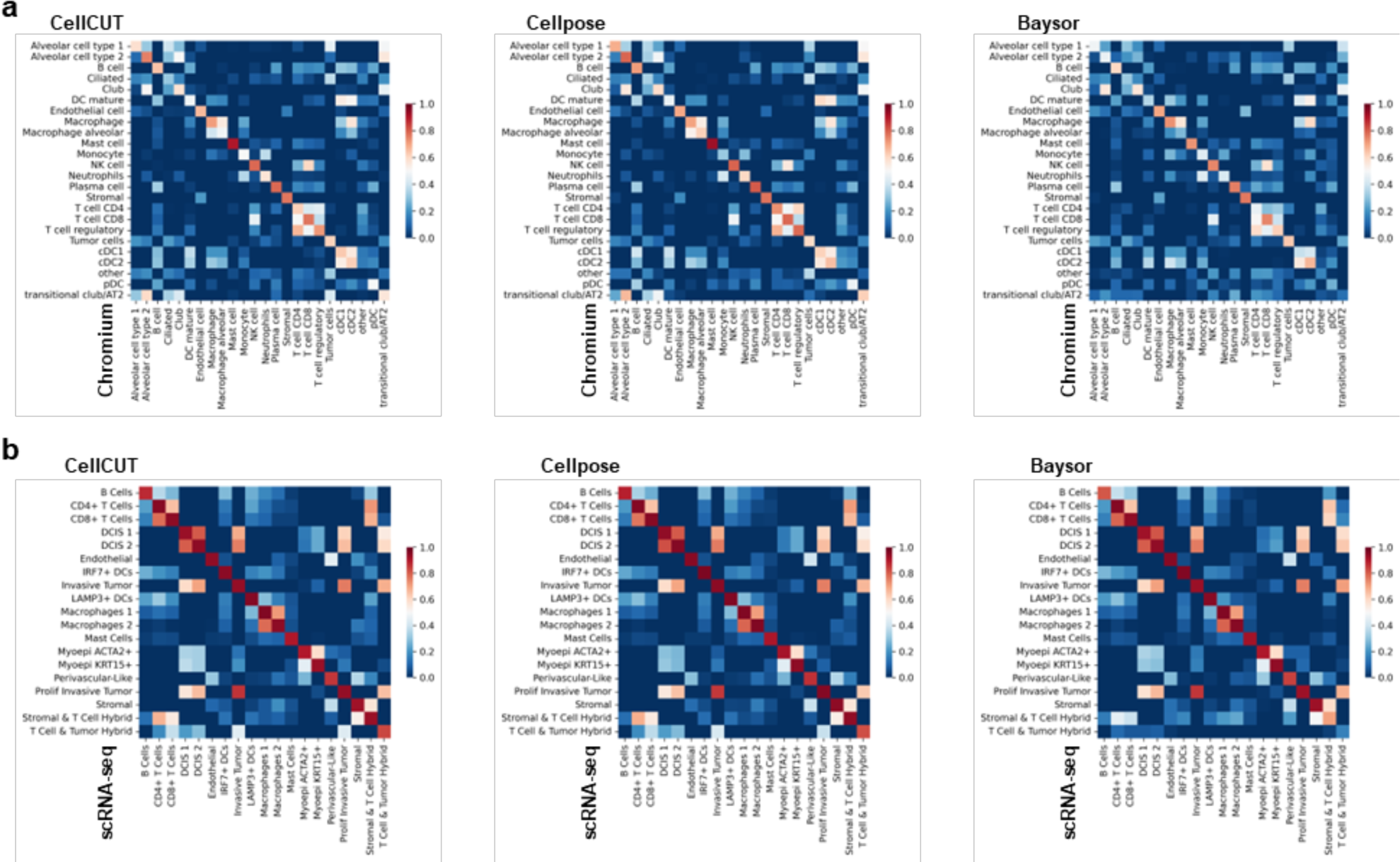
Correlation heatmaps of average expression between spatial data (y-axis) and scRNA-seq data (x-axis). **a**, Correlation heatmaps in the NanoString NSCLC dataset. **b**, Correlation heatmaps in the Xenium breast cancer dataset.

**Supplementary Fig. S13.**
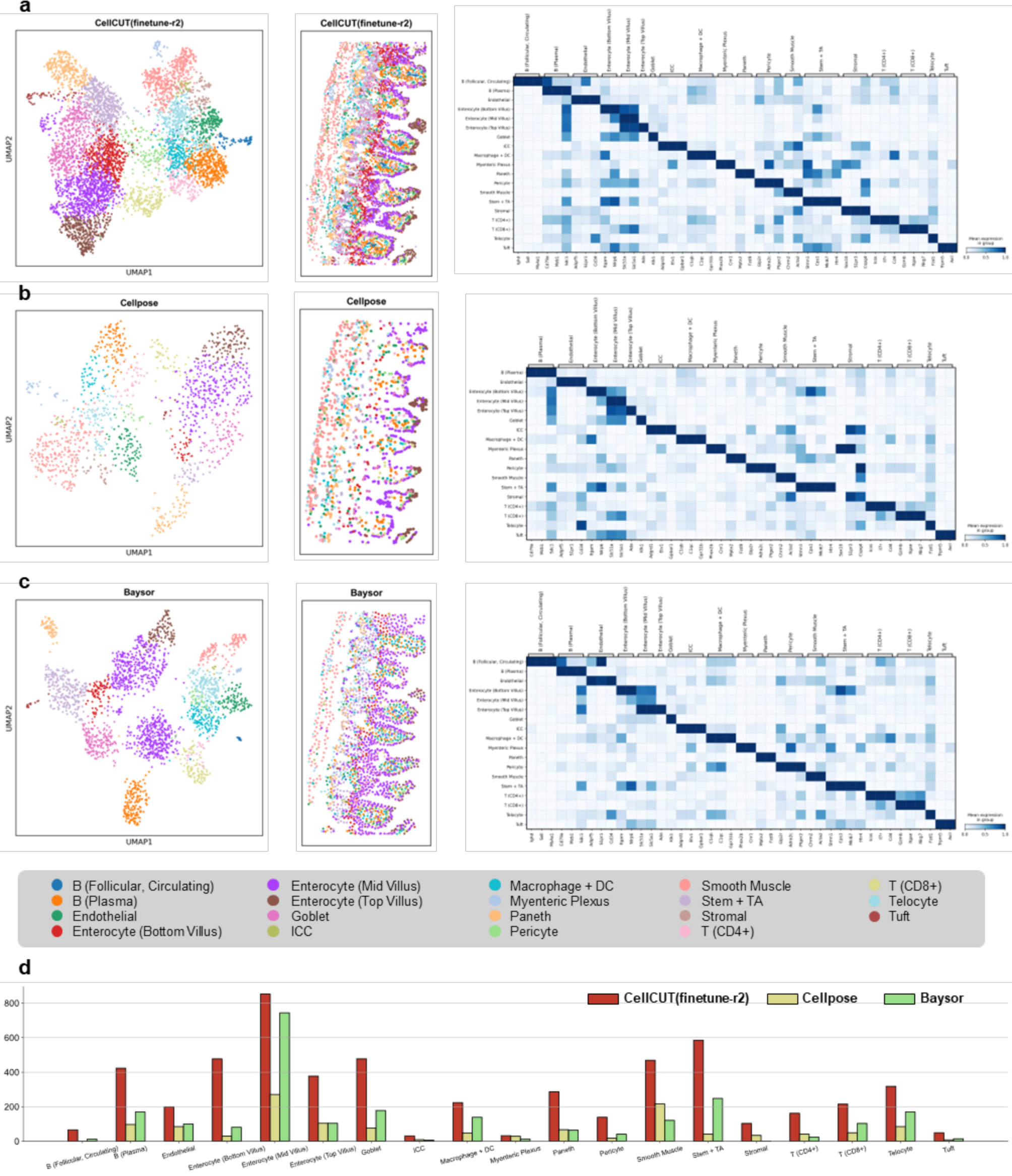
Comparisons of cell types in the MERFISH mouse ileum dataset recovered from different methods. **a**, **b**, and **c** are the cell type annotations produced by CellCUT, Cellpose, and Baysor, respectively. From left to right are UMAP of all identified cells colored based on Leiden clusters, cell type spatial distributions, and marker gene expressions by each of the identified cell types, respectively. **d**, The number of each cell type identified by each of the segmentation methods in a-c.

**Supplementary Fig. S14.**
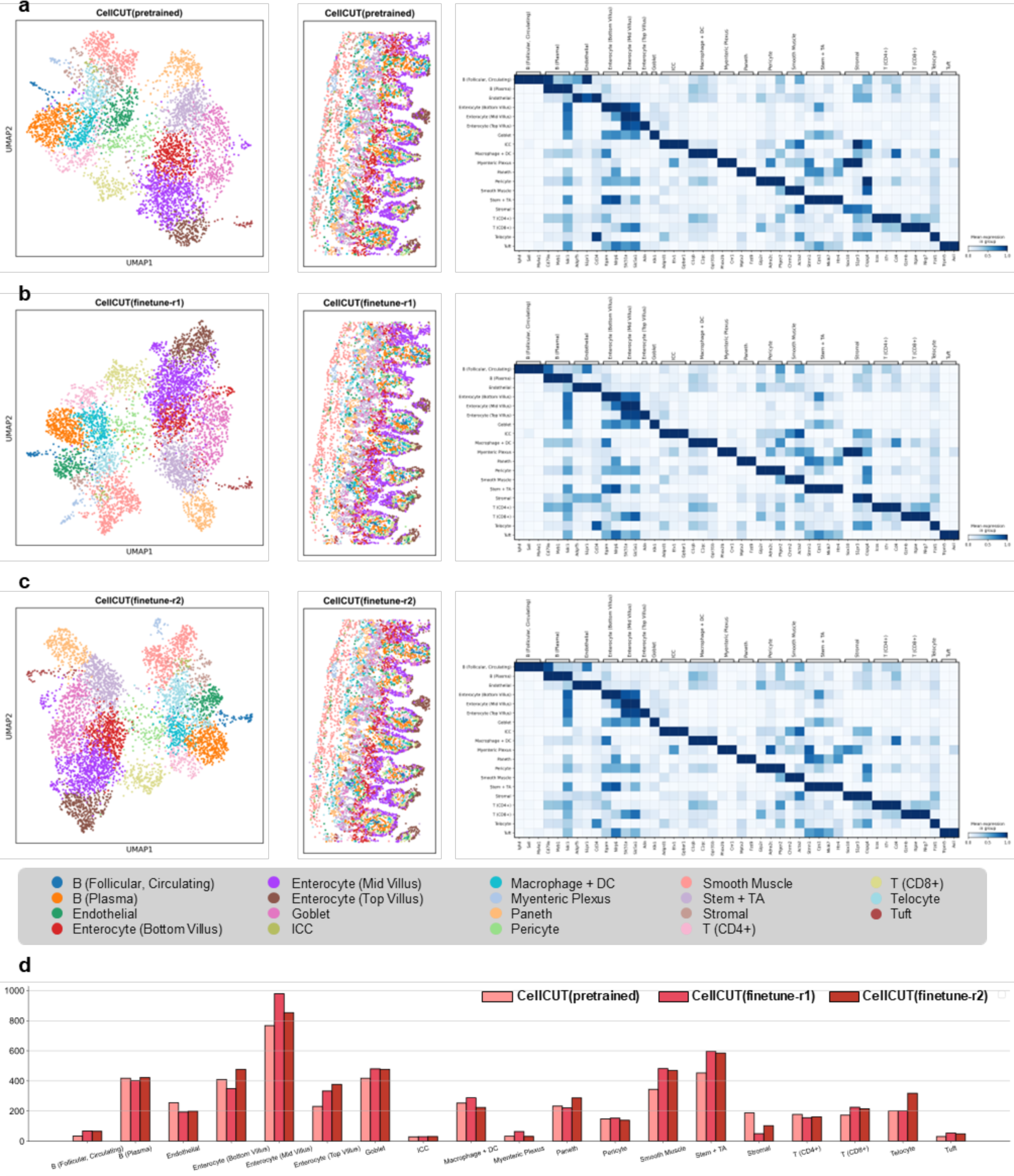
Comparisons of cell types in the MERFISH mouse ileum dataset recovered from different methods. **a**, **b**, and **c** are the cell type annotations produced by CellCUT(pretrained), CellCUT(fineture-r1), and CellCUT(fineture-r2), respectively. From left to right are UMAP of all identified cells colored based on Leiden clusters, cell type spatial distributions, and marker gene expressions by each of the identified cell types, respectively. **d**, The number of each cell type identified by each of the segmentation methods in a-c.

**Supplementary Fig. S15.**
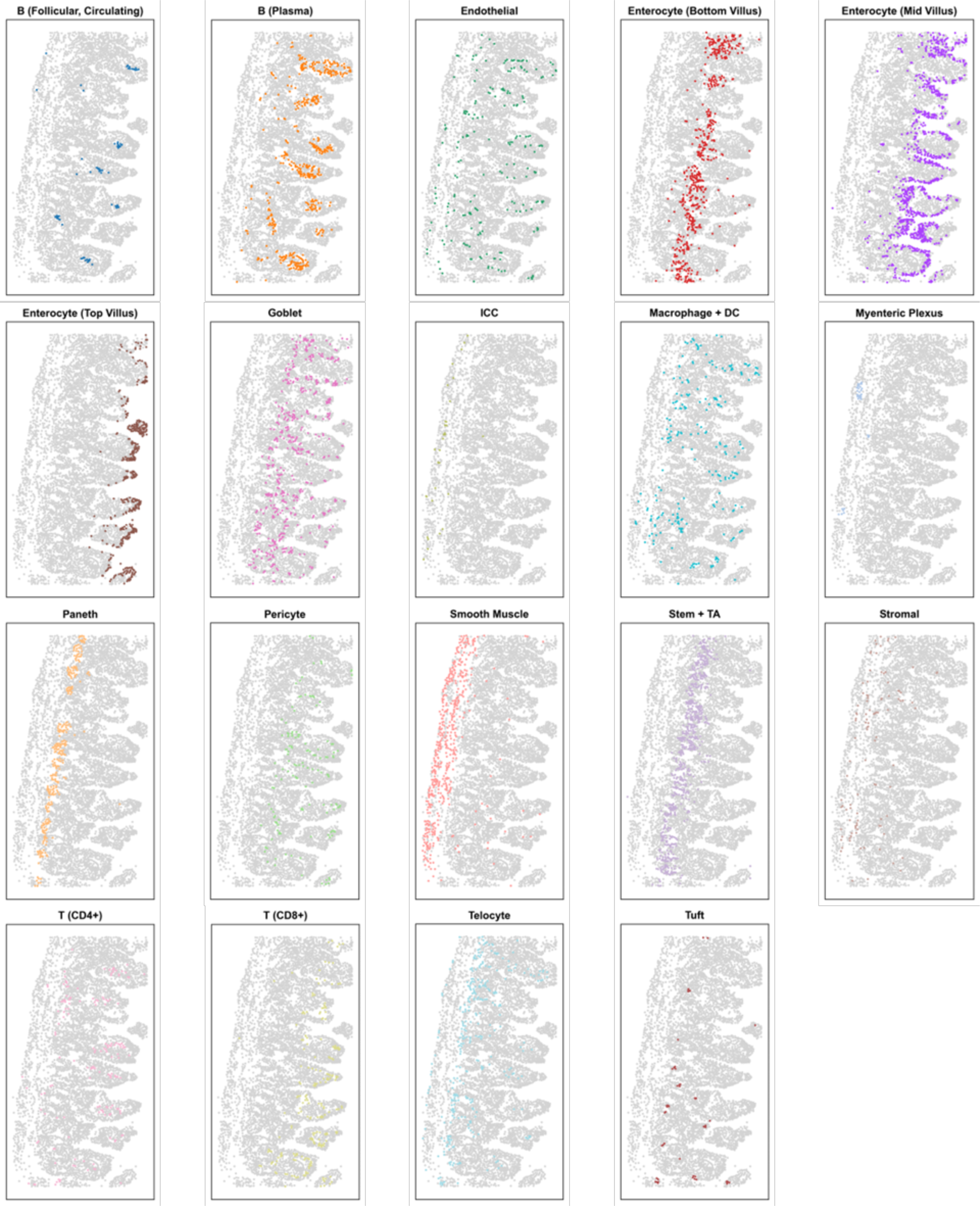
Spatial distributions of all cell types identified by CellCUT(fineture-r2) in the MERFISH mouse ileum dataset. The gray dots represent the location of all cells and the colored dots represent the location of the indicated cell type.

**Supplementary Fig. S16.**
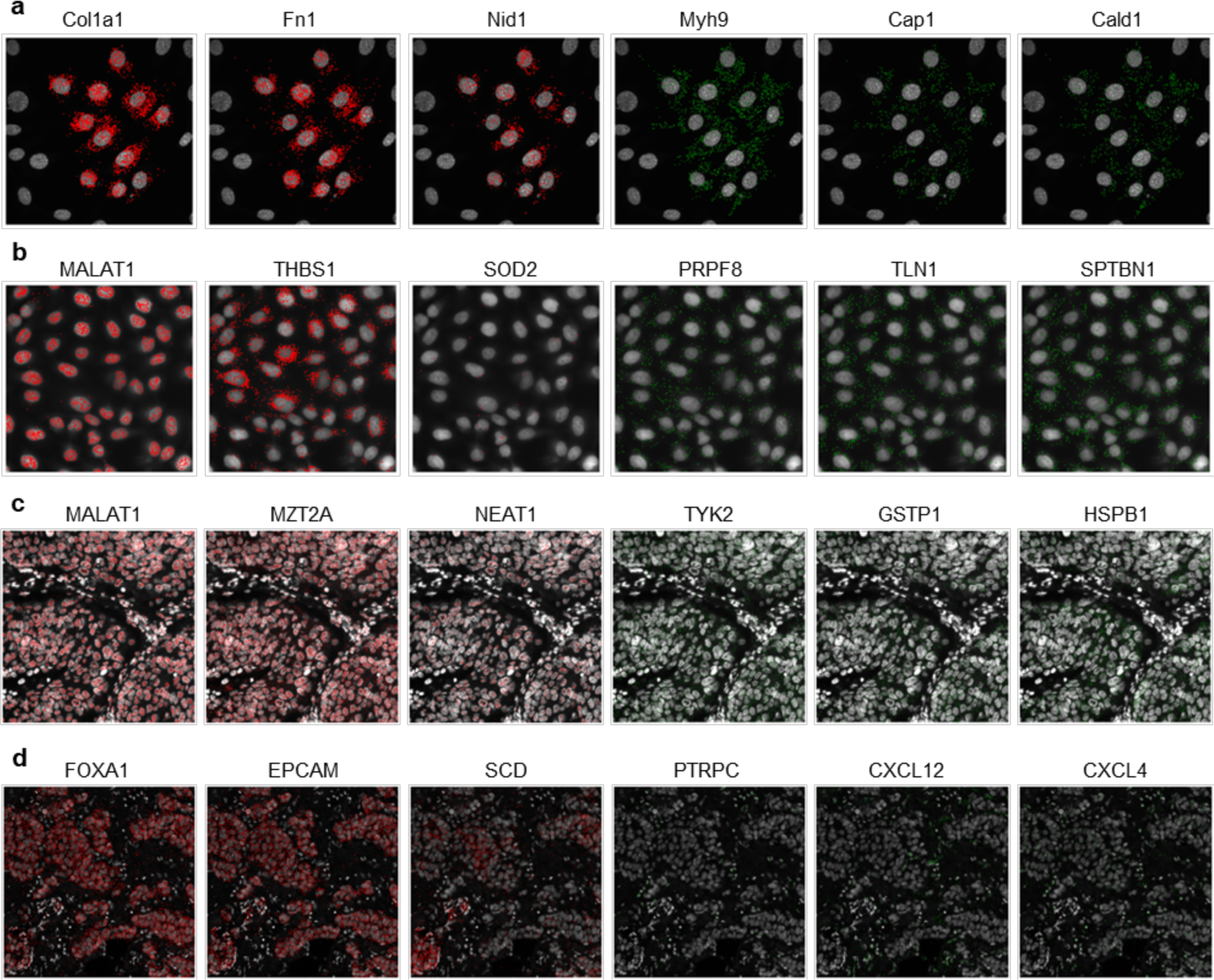
Visualization of spatial distribution of genes. **a**, **b**, **c**, and **d** show selected visualization results of tagging the top three genes with the most significant differences for each category in the SeqFISH+ NIH/3T3 dataset, MERFISH u2os dataset, NanoString NSCLC dataset, and Xenium breast cancer dataset, respectively. The red dots denote transcripts enriched in the nucleus, and the green dots denote transcripts enriched in the cytoplasm.

## References

1. Marx, V. Method of the Year: spatially resolved transcriptomics. Nat Methods 18, 9–14 (2021).

2. Shi, Y. et al. Decoding the spatiotemporal regulation of transcription factors during human spinal cord development. Cell Res (2024) doi:10.1038/s41422-023-00897-x.

3. Williams, C. G., Lee, H. J., Asatsuma, T., Vento-Tormo, R. & Haque, A. An introduction to spatial transcriptomics for biomedical research. Genome Med 14, 68 (2022).

4. Wang, X. et al. Three-dimensional intact-tissue sequencing of single-cell transcriptional states. Science 361, eaat5691 (2018).

5. He, S. et al. High-plex imaging of RNA and proteins at subcellular resolution in fixed tissue by spatial molecular imaging. Nat Biotechnol 40, 1794–1806 (2022).

6. Janesick, A. et al. High resolution mapping of the tumor microenvironment using integrated single-cell, spatial and in situ analysis. Nat Commun 14, 8353 (2023).

7. Moffitt, J. R. et al. High-throughput single-cell gene-expression profiling with multiplexed error-robust fluorescence in situ hybridization. Proc. Natl. Acad. Sci. U.S.A. 113, 11046–11051 (2016).

8. Lubeck, E., Coskun, A. F., Zhiyentayev, T., Ahmad, M. & Cai, L. Single-cell in situ RNA profiling by sequential hybridization. Nat Methods 11, 360–361 (2014).

9. Eng, C.-H. L. et al. Transcriptome-scale super-resolved imaging in tissues by RNA seqFISH+. Nature 568, 235–239 (2019).

10. Bressan, D., Battistoni, G. & Hannon, G. J. The dawn of spatial omics. Science 381, eabq4964 (2023).

11. Moses, L. & Pachter, L. Museum of spatial transcriptomics. Nat Methods 19, 534–546 (2022).

12. Qiu, X., et al. Spateo: Multidimensional Spatiotemporal Modeling of Single-Cell Spatial Transcriptomics. http://biorxiv.org/lookup/doi/10.1101/2022.12.07.519417(2022) doi:10.1101/2022.12.07.519417.

13. Stringer, C., Wang, T., Michaelos, M. & Pachitariu, M. Cellpose: a generalist algorithm for cellular segmentation. Nat Methods 18, 100–106 (2021).

14. Schmidt, U., Weigert, M., Broaddus, C. & Myers, G. Cell Detection with Star-Convex Polygons. in Medical Image Computing and Computer Assisted Intervention – MICCAI 2018 (eds. Frangi, A. F., Schnabel, J. A., Davatzikos, C., Alberola-López, C. & Fichtinger, G.) vol. 11071 265–273 (Springer International Publishing, Cham, 2018).

15. Greenwald, N. F. et al. Whole-cell segmentation of tissue images with human-level performance using large-scale data annotation and deep learning. Nat Biotechnol 40, 555–565 (2022).

16. Wang, Y. et al. GeneSegNet: a deep learning framework for cell segmentation by integrating gene expression and imaging. Genome Biol 24, 235 (2023).

17. Littman, R. et al. Joint cell segmentation and cell type annotation for spatial transcriptomics. Molecular Systems Biology 17, e10108 (2021).

18. Qian, X. et al. Probabilistic cell typing enables fine mapping of closely related cell types in situ. Nat Methods 17, 101–106 (2020).

19. He, Y. et al. ClusterMap for multi-scale clustering analysis of spatial gene expression. Nat Commun 12, 5909 (2021).

20. Petukhov, V. et al. Cell segmentation in imaging-based spatial transcriptomics. Nat Biotechnol 40, 345–354 (2022).

21. Chen, H., Li, D. & Bar-Joseph, Z. SCS: cell segmentation for high-resolution spatial transcriptomics. Nat Methods 20, 1237–1243 (2023).

22. Jin, K. et al. Bering: *Joint Cell Segmentation and Annotation for Spatial Transcriptomics with Transferred Graph Embeddings*. http://biorxiv.org/lookup/doi/10.1101/2023.09.19.558548 (2023) doi:10.1101/2023.09.19.558548.

23. Fu, X., et al. Biologically-Informed Self-Supervised Learning for Segmentation of Subcellular Spatial Transcriptomics Data. http://biorxiv.org/lookup/doi/10.1101/2023.06.13.544733 (2023) doi:10.1101/2023.06.13.544733.

24. Zhang, Y. et al. Reference-based cell type matching of in situ image-based spatial transcriptomics data on primary visual cortex of mouse brain. Sci Rep 13, 9567 (2023).

25. Chen, A. et al. Spatiotemporal transcriptomic atlas of mouse organogenesis using DNA nanoball-patterned arrays. Cell 185, 1777–1792.e21 (2022).

26. Mah, C. K., et al. Bento: A Toolkit for Subcellular Analysis of Spatial Transcriptomics Data. http://biorxiv.org/lookup/doi/10.1101/2022.06.10.495510 (2022) doi:10.1101/2022.06.10.495510.

27. Ronneberger, O., Fischer, P. & Brox, T. U-Net: Convolutional Networks for Biomedical Image Segmentation. in Medical Image Computing and Computer-Assisted Intervention – MICCAI 2015 (eds. Navab, N., Hornegger, J., Wells, W. M. & Frangi, A. F.) vol. 9351 234–241 (Springer International Publishing, Cham, 2015).

28. He, K., Zhang, X., Ren, S. & Sun, J. Deep Residual Learning for Image Recognition. in 2016 IEEE Conference on Computer Vision and Pattern Recognition (CVPR) 770–778 (IEEE, Las Vegas, NV, USA, 2016). doi:10.1109/CVPR.2016.90.

29. Turaga, S. C. et al. Convolutional Networks Can Learn to Generate Affinity Graphs for Image Segmentation. Neural Computation 22, 511–538 (2010).

30. De Brabandere, B., Neven, D. & Van Gool, L. Semantic Instance Segmentation with a Discriminative Loss Function. Preprint at http://arxiv.org/abs/1708.02551 (2017).

31. Lin, Z., Wei, D., Lichtman, J. & Pfister, H. PyTorch Connectomics: A Scalable and Flexible Segmentation Framework for EM Connectomics. Preprint at http://arxiv.org/abs/2112.05754 (2021).

32. Caicedo, J. C. et al. Nucleus segmentation across imaging experiments: the 2018 Data Science Bowl. Nat Methods 16, 1247–1253 (2019).

33. Velickovic, P., et al. GRAPH ATTENTION NETWORKS. (2018).

34. Ding, K., Xu, Z., Tong, H. & Liu, H. Data Augmentation for Deep Graph Learning: A Survey. SIGKDD Explor. Newsl. 24, 61–77 (2022).

35. Beier, T. et al. Multicut brings automated neurite segmentation closer to human performance. Nat Methods 14, 101–102 (2017).

36. Chopra, S. The partition problem.

37. Hong, B. et al. Joint reconstruction of neuron and ultrastructure via connectivity consensus in electron microscope volumes. BMC Bioinformatics 23, 453 (2022).

38. Keuper, M. et al. Efficient Decomposition of Image and Mesh Graphs by Lifted Multicuts. in 2015 IEEE International Conference on Computer Vision (ICCV) 1751–1759 (IEEE, Santiago, Chile, 2015). doi:10.1109/ICCV.2015.204.

39. Wolf, F. A., Angerer, P. & Theis, F. J. SCANPY: large-scale single-cell gene expression data analysis. Genome Biol 19, 15 (2018).

40. Cisar, C., Keener, N., Ruffalo, M. & Paten, B. A unified pipeline for FISH spatial transcriptomics. Cell Genomics 3, 100384 (2023).

